# Predicting the invasiveness of threshold-dependent gene drives

**DOI:** 10.1101/2025.11.20.689598

**Authors:** Isabel K. Kim, Philipp W. Messer

## Abstract

Gene drives hold great promise for controlling disease vectors or invasive species due to their capacity to rapidly spread through a population from a small initial release. This same property also raises serious concerns about unintended spillover into non-target populations. Threshold-dependent gene drive systems, which can spread only when introduced above a critical population frequency, have been proposed as a more controllable alternative, yet their invasion dynamics in spatially structured populations remain poorly understood. Here, we analyze invasion criteria for threshold-dependent gene drives in continuous-space populations using deterministic reaction-diffusion models and individual-based simulations that better capture the stochasticity of real-world populations. We find substantial variability in invasion outcomes in the individual-based models. Low-threshold modification drives with small fitness costs frequently spread across a wide range of release sizes, including introductions far below those required to succeed in diffusion models. In contrast, threshold-dependent suppression drives exhibit qualitatively different behavior: stochastic effects at low density can often disrupt wavefronts or produce persistent chasing cycles, generally reducing invasion success relative to diffusion-model expectations. Overall, our results show that the spatial containment of threshold-dependent gene drives is more complex than predicted by non-spatial or purely deterministic models, highlighting the importance of spatially explicit analyses when evaluating their real-world performance.

## 1 Introduction

The idea of designing self-propagating genetic constructs that can spread specific genetic changes through a population has fascinated scientists for a long time [1–3]. With the advent of CRISPR technology, the engineering of such “gene drives” has finally come within reach. Over recent years, successful drive designs have been demonstrated in microbes [4, 5], plants [6, 7], insects [8–12], and mammals [13, 14]. Their potential applications are manifold; for instance, a gene drive could be designed to spread a payload gene in mosquitoes that prevents them from transmitting malaria parasites [15], or to revert a recently acquired insecticide resistance mutation to the wild type [16]. Beyond these so-called population “modification” drives, a “suppression” drive can directly deplete or even eradicate a target population when it biases the population’s sex ratio [11] or causes recessive sterility [9]. Suppression drives could provide new means to combat disease-carrying mosquito vectors [17], invasive species [18], and agricultural pests [19].

The key feature of a gene drive—its ability to spread quickly throughout an entire population from a small release—also underlies the major concern against their use, as it limits our ability to control and confine a release. If a gene drive spreads into areas outside of the target region, this could result in the unintended modification or extinction of a non-target population, with potentially severe ecological, economic, and political consequences [20, 21].

Threshold-dependent drive systems could provide a solution to this problem because they are only expected to spread when introduced above a given threshold frequency in the population [22, 23]. One classic example is an underdominance system, in which heterozygotes are less fit than either homozygote [24–26]. Davis et al. [24] proposed an engineered underdominance system involving two constructs at two genetically unlinked loci, where each construct would contain a lethal element and a suppressor of the other construct’s lethal element. Thus, only individuals who inherit neither or both constructs would survive. A desirable gene could be attached to one of the constructs and would increase in frequency as the underdominance system spreads.

More recently, toxin-antidote gene drives have been proposed that use CRISPR to disrupt an essential gene (for the “toxin” effect) while at the same time providing a recoded copy of the target gene within the drive construct that cannot be cut (for the “antidote” effect) [27, 28]. These drives spread by converting wild-type alleles to disrupted alleles, which can be rescued only when the individual also inherits a drive allele. The invasion threshold of such a system can be tuned through the fitness costs associated with the different alleles. CRISPR-based toxin-antidote gene drive systems have already been demonstrated in insects [29–32] and plants [6, 7]. Other forms of threshold-dependent gene drives include *Wolbachia* [33, 34], Medea [35, 36], and reciprocal chromosomal translocations [1, 37].

In principle, a threshold-dependent gene drive can remain confined to a target population as long as the drive never locally exceeds its threshold frequency. Several modeling studies have explored the conditions for this in idealized models where a target deme is connected to a non-target deme by migration, finding that invasion can indeed be prevented as long as the migration rate between demes is low enough [20, 38–41]. A key assumption of these models is that the individual demes are panmictic (i.e., there is no structure within each deme, so that all individuals have an equal chance of mating with each other).

Real-world populations, by contrast, are usually far away from this panmixia assumption, especially when they inhabit larger geographic ranges where individuals tend to mate with those born nearby. This makes it difficult to interpret the threshold frequency required for drive invasion. Champer et al. [42] pointed out how the panmixia assumption would roughly translate to migrants being placed at random positions within the geographic range of the non-target population. Yet if migrants arrive spatially clustered within their new deme—such as around a ship harbor—the drive may invade much more readily. In this case, it could be introduced at a low overall frequency in the population, but at a high enough local frequency around the release site to spread outwards and eventually through the entire population [42]. These findings highlight the challenges of predicting how threshold-dependent gene drives will spread in realistic spatial populations, which poses a key concern for current safety assessments.

Reaction-diffusion models could help us address this challenge by providing insights into the factors that determine the invasiveness of threshold-dependent gene drives in continuous-space populations. Barton and Turelli [43] used this framework to study the spread of a simple underdominance allele, deriving the so-called “critical bubble”—a unique frequency distribution at which the effects of diffusion and selection exactly balance each other out. If introduction frequencies are everywhere above the critical bubble, the allele spreads, and if introduction frequencies are everywhere below the critical bubble, the allele is lost—making the critical bubble the spatial analogue to the threshold frequency of panmictic models. Tanaka et al. [44] contributed significant insights toward adapting this approach to gene drive systems.

However, the reaction-diffusion framework underlying the critical bubble concept rests on several assumptions that are often strongly violated in real populations, potentially limiting its applicability. The model assumes, for example, both weak selection and an infinitely large, one-dimensional (1D) habitat in which individuals disperse over distances that are small relative to the population’s overall range [45]. Moreover, populations are not treated as discrete individuals but as a continuous function of local allele frequencies, which changes deterministically over time and space. In real populations, by contrast, the discrete nature of individuals results in stochasticity that can play a critical role [46]. For instance, genetic drift can lead to the loss of alleles that would always be predicted to spread in a deterministic model, and phenomena such as “chasing” dynamics [47] can profoundly alter the behavior of a drive. The reaction–diffusion model used to derive the critical bubble also assumes a constant population density across the range, effectively invoking soft selection, where fitness influences allele frequencies but not population size [48]. Consequently, such models cannot adequately describe suppression drives that directly reduce local population densities in the regions they invade. Perhaps most importantly, migrants carrying the drive into a non-target population are unlikely to arrive in the distinctive spatial pattern that defines the critical bubble. Instead, their initial distribution may exceed the threshold in some areas while falling below it in others, undermining the predictive value of the critical bubble concept for invasion outcomes. Finally, no equivalent of the critical bubble has yet been derived for 2D populations, which likely encompass the majority of natural systems that inhabit large geographic ranges.

In this study, we devise individual-based simulation models of threshold-dependent gene drive systems in spatial populations to comprehensively evaluate their invasiveness under more realistic assumptions. Using this framework, we first test the concept of a critical bubble for a simple underdominance system and find that it generally lies within the range of release conditions above which the allele can spread. However, invasion can exhibit substantial stochasticity; a system can often spread from releases well below the predicted critical bubble or fail to spread despite being released well above it. We then examine more realistic scenarios where a drive is introduced at a constant frequency over a local region. In this case, invasion can occur from releases much smaller than the critical bubble, yet we show that numerical reaction–diffusion predictions can still be quite accurate. We further extend this framework to homing underdominance gene drives as well as toxin–antidote modification and suppression drives, presenting results for both one- and two-dimensional populations. For suppression drives, we observe qualtatively different outcomes that deterministic reaction–diffusion models fail to capture entirely. Throughout, we identify parameter regimes in which reaction–diffusion predictions mostly hold and where they break down.

## 2 The critical bubble

Barton and Turelli [43] developed a reaction-diffusion model of a single-locus underdominance system in 1D continuous space in order to derive the unique frequency distribution at which the effects of selection and diffusion exactly cancel, which they called the critical bubble. In their model, homozygotes for the underdominance allele (*d*) have fitness *f*_*dd*_ = 1 + 2*s*_*d*_, homozygotes for the wild-type allele (*w*) have fitness *f*_*ww*_ = 1, and heterozygotes have fitness *f*_*dw*_ = 1 + *s*_*d*_ − *s*_*u*_. Let *p*(*t*) denote the overall frequency of the underdominance allele in the population in generation *t*. Under panmixia, and assuming that selection is weak, the expected change in frequency per generation is 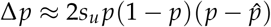, where 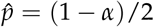 specifies the panmictic invasion threshold (obtained by setting *p* = 0) and *α* = *s*_*d*_/*s*_*u*_ is the ratio of directional over underdomi-nance selection. A necessary condition for spread is therefore that *s*_*d*_ > 0 with 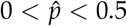.

In a reaction-diffusion model, the frequency of the underdominance allele becomes a function not only of time *t* but also of spatial position *x*. The spatiotemporal dynamics are then described by the combined effects of a diffusion term, which models the change in local allele frequencies due to random dispersal, and a reaction term, which models the local change due to selection. Barton and Turelli [43] applied this approach to their underdominance system in a 1D continuous-space population, obtaining a reaction-diffusion equation of the form

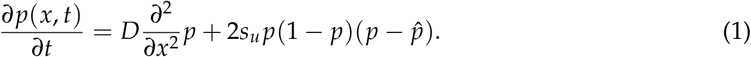

The diffusion parameter *D* = *σ*^2^/2 is defined here by the variance in displacement distance (*σ*^2^) between mother and offspring, assuming a symmetric dispersal kernel that is not too heavy-tailed. For a list of all variables used in our paper, see Table S1.

In Text S1, we recapitulate the detailed derivation of the critical bubble distribution from this equation. Here, we just summarize two of its key properties. The first is the frequency at the center of the critical bubble, which specifies the maximum height of the curve and is given by

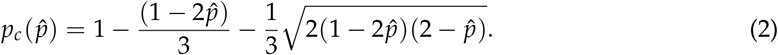

The second is the area under the critical bubble, which is given by

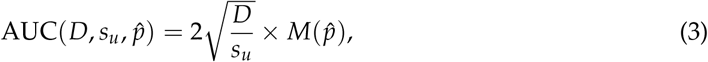

where

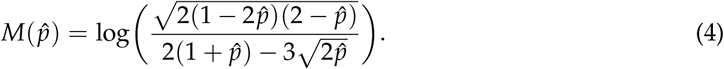

Note that the factor 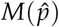 depends only on the system’s panmictic invasion threshold 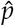. The critical release size under this distribution will be proportional to the area under the curve. For example, if the population density is 100,000 individuals per unit length, the critical release size would be 100,000 × AUC.

The shape of the critical bubble effectively depends on only two parameters: (i) the ratio *D*/*s*_*u*_ of the diffusion coefficient to the underdominance selection coefficient, and (ii) the panmictic invasion threshold 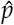. Figure 1 illustrates how each of these parameters influences the shape. As *D*/*s*_*u*_ increases, the critical bubble becomes wider, with the area under the curve scaling linearly with 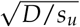 (Figure 1A). The panmictic threshold 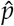 determines the height of the curve (*p*_*c*_) through Eq. 2. As 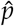 increases, the critical bubble must grow to provide local frequencies that exceed the threshold by a sufficient amount (Figure 1B and D). This results in an almost linear relationship between 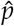 and the area under the critical bubble (Figure 1C).

**Figure 1:**
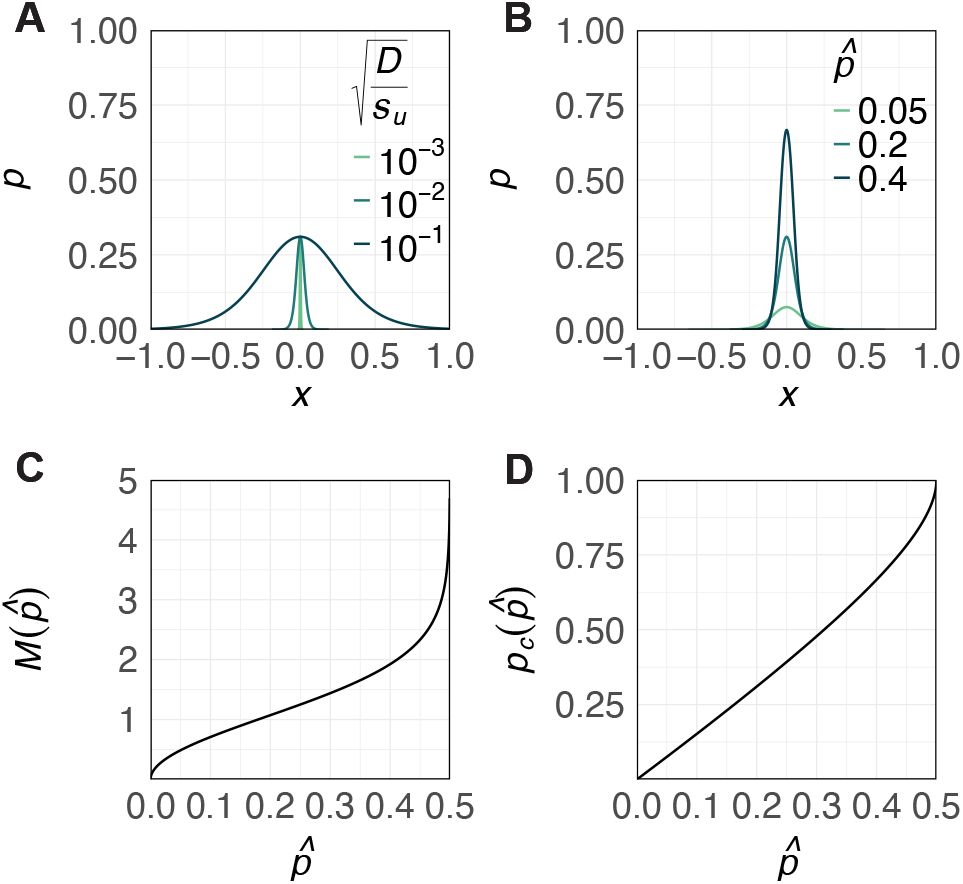
Variables influencing the shape of the critical bubble. **A**. The effect of 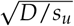 on the width of the critical bubble, where *D* represents the diffusion constant and *s*_*u*_ represents the underdominance selection coefficient. **B**. The effect of the invasion threshold 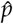 on the height of the critical bubble. **C**. The relationship between the invasion threshold 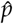 and 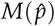 (Eq. 4), which is proportional to the area under the critical bubble (Eq. 3). **D**. The relationship between the invasion threshold 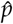 and the height of the critical bubble *p*_*c*_ (Eq. 2).

## 3 Individual-based simulation model in SLiM

To test how well these mathematical predictions from diffusion theory apply to populations composed of discrete individuals, we created an individual-based simulation model of a diploid, sexually reproducing population in SLiM version 5.0 [49]. In our initial model, individuals inhabit a 1D continuous line of 1-unit length with toroidal boundaries and a global carrying capacity of 100,000 individuals. Simulations begin with wild-type individuals of random sex randomly distributed along the line. In an effort to resemble the reaction-diffusion model (where all processes are completely local) as closely as possible, females always mate with their nearest male neighbor. In rare cases where a female cannot find a mate (within a width containing 100 individuals at equilibrium), she does not reproduce. To maintain roughly constant densities across space, individuals overproduce offspring, with litter sizes drawn from a Poisson distribution with a mean of 15 and sex assigned at random. Discrete generations are obtained by removing parents after mating.

Offspring then experience viability selection and density-dependent competition. In Barton and Turelli [43]’s simple underdominance model, homozygotes for the underdominance allele have the highest fitness of *f*_*dd*_ = 1 + 2*s*_*d*_. To translate these fitness values into survival probabilities (*v*) while maintaining their ratios, we scale all fitness values by the maximum fitness such that *v*_*dd*_ = *f*_*dd*_/ *f*_*dd*_ = 1, *v*_*dw*_ = *f*_*dw*_/ *f*_*dd*_, and *v*_*ww*_ = *f*_*ww*_/ *f*_*dd*_. After viability selection, surviving offspring are placed at their mother’s position and compete locally for resources. We implement competition using a resource-explicit model introduced by Champer et al. [50] with a global carrying capacity of 100,000 individuals, distributed over 2500 local resource nodes that are equidistantly spaced over the arena line. Individuals can only forage for resources in their local neighborhood, such that success will be inversely proportional to the number of local competitors. After foraging has taken place, the population reaches capacity, with an approximately constant density across the arena line. The rationale for choosing this model was again to closely resemble diffusion theory assumptions of constant local population density and soft selection.

Surviving offspring after density competition are displaced from their mother’s position by a random distance drawn from a Gaussian distribution with mean zero and variance *σ*^2^ = 2*D*, esulting in an average dispersal distance of 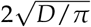. Diffusion theory assumes an infinite arena in which individuals move short distances. Thus, we expect model outcomes to align more closely with reaction-diffusion predictions at low values of *D*, while behavior at high *D* should more closely align with panmictic predictions.

We allowed the wild-type population to equilibrate for 10 generations before introducing homozygotes for the underdominance allele in a given release profile. Simulations continued until the underdominance allele reached fixation or loss, or until 100 generations had passed since the introduction.

## 4 Testing the critical bubble in SLiM

In diffusion models, space is assumed to be infinite, but in our simulation, the arena is bounded between *x* = 0 and *x* = 1. For critical bubble releases, we only modeled the portion of the bubble that falls within the arena bounds. For underdominance systems with high selection coefficients (*s*_*u*_) or in populations with low diffusion constants (*D*) (e.g., Figure S1A), most of the critical bubble is captured within these boundaries, while for low *s*_*u*_ or high *D*, bubbles can extend far beyond the boundaries (e.g., Figure S1B). We centered the bubble at *x* = 0.5 and converted each individual to a homozygote for the underdominance allele with probability equal to the corresponding critical bubble frequency at its position *x*. This method, on average, produces a spatial frequency distribution of the underdominance allele equal to the critical bubble.

To test whether releases everywhere above the critical bubble lead to fixation and releases everywhere below the critical bubble lead to loss, we applied a multiplicative factor *ϕ* to critical bubble frequencies such that a value of *ϕ* = 1 produces a distribution equal to the critical bubble, *ϕ* > 1 produces a distribution above the critical bubble, and *ϕ* < 1 produces a distribution below the critical bubble. We varied *ϕ* between 0.5 and 2, with 20 replicates per value, and recorded the average release size under the given frequency distribution as well as the fraction of replicates in which the underdominance allele increased in frequency, denoted by *P*(increase).

Figure 2A and B show *P*(increase) for two different underdominance systems as a function of the average total release size, obtained by varying *ϕ*. Both underdominance systems have panmictic invasion thresholds of 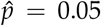, but releases with global frequencies far below 5% were sufficient for the allele to invade (a release size of 5000 lies outside of the range of the x-axis shown in the figure). Using the drc package in R [51], we fit a logistic curve to each plot and defined the SLiM critical release size to be the release size at which *P*(increase) = 0.5. The SLiM critical release size was close to the predicted critical bubble release size of 100,000 × AUC when the bubble could be well approximated in our simulations (Figure 2C). However, outcomes around the critical release size varied substantially due to the stochasticity in our simulations: sometimes, the underdominance allele could spread when released well below the critical bubble, and sometimes, the allele could be introduced far above the critical bubble yet still decline (Figure 2A and B). To measure this variability for a given underdominance system, we defined the “transition range” to be release sizes that produced 0.05 ≤ *P*(increase) ≤ 0.95.

**Figure 2:**
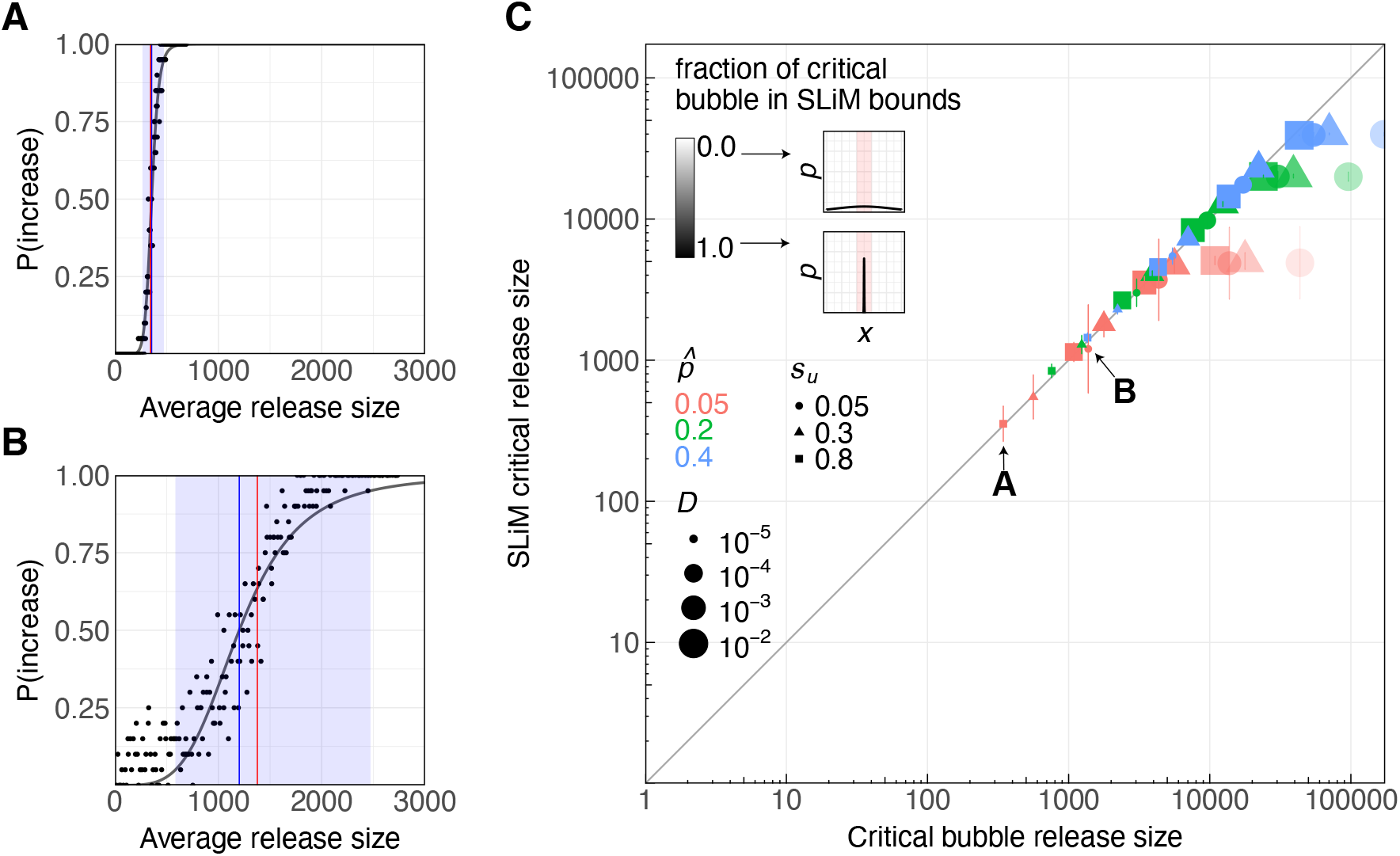
Invasion probability under critical bubble releases. **A**. Spread of an underdominance system with *D* = 10^−5^, *s*_*u*_ = 0.8, and 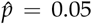. Each dot reflects 20 simulation runs at a given multiplicative factor *ϕ* applied to the critical bubble. The average release size for a given *ϕ* is shown on the x-axis and the fraction of replicates in which the underdominance system increased in frequency is shown on the y-axis, denoted *P*(increase). We fit a logistic curve (shown in grey) to each plot using the drc package in R [51]. The shaded blue region represents the transition range where the underdominance allele has 0.05 ≤ *P*(increase) ≤ 0.95, and the blue line marks the SLiM critical release size at which *P*(increase) = 0.5 based on the fitted curve. The red line denotes the predicted threshold release size under the full critical bubble distribution: 100,000 × AUC (corresponding to *ϕ* = 1). **B**. Same as **A**, but for an underdominance system with *D* = 10^−5^, *s*_*u*_ = 0.05, and 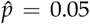. **C**. Comparison of the SLiM critical release size to the critical bubble release size. Each point reflects a different underdominance system, with bars representing its transition range in SLiM. The transparency of the point reflects the portion of the underdominance system’s critical bubble that is contained within the SLiM bounds. The grey line denotes *y* = *x*.

In general, the transition range widens with smaller values of *s*_*u*_ when holding other parameters constant. For the underdominance system shown in Figure 2B, for example, outcomes were so variable that a release of only a handful of underdominance-allele homozygotes was enough for the allele to spread in a notable fraction of runs. This result is consistent with Jansen et al. [52], who found that in a finite population, a weakly deleterious *Wolbachia* system initially present in only one individual could spread to fixation at rates similar to that of a zero-threshold system. This is because when the local population density is finite, genetic drift can allow a threshold-dependent system to rise well above 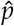 in a local area and spread. This is easier at low 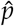 and for small *s*_*u*_, and in populations with lower density. With increasing population density, we would expect the transition range to contract, and in the limit of an infinite population density (as in diffusion models), invasion would become deterministic.

## 5 Rectangular releases

Since accidental introductions into a non-target population are unlikely to occur in the peculiar critical bubble shape, we next considered a simpler and more generic introduction scenario: a concentrated release of the underdominance allele at 100% frequency (*p*_0_) within a region of set width. To simulate this “rectangular release” in SLiM, we replaced all individuals located within the given release width (i.e, segment of the arena line) with homozygotes for the underdominance allele, ran the simulation 20 times, and recorded the fraction of replicates in which the allele increased in frequency and the average release size under this distribution (which, on average, equals the carrying capacity of 100,000 times the release width). For a given underdominance system, we varied the release width and found the SLiM critical release size and the transition range using the same logistic-curve-fitting protocol described in Section 4.

Figure 3C compares the critical rectangular release size in SLiM to the critical bubble release size for a range of underdominance systems. For systems with low diffusion constants *D*, high selection coefficients *s*_*u*_, and high thresholds 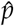, as in Figure 3B, the critical bubble is tall and narrow. A rectangle with height *p*_0_ = 1 and equal area to this critical bubble resembles it more closely than does a system with high *D*, low *s*_*u*_, or low 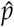, with a shorter and wider bubble (as in Figure 3A). Thus, it is not surprising that underdominance systems with tall, narrower bubbles had critical rectangular release sizes in SLiM that were more similar to their critical bubble release sizes. However, across all systems tested, the critical rectangular release size in SLiM was consistently less than the critical bubble release size. In other words, underdominance alleles could always spread more easily when introduced at a 100% frequency over a critical width than from the critical bubble release shape.

**Figure 3:**
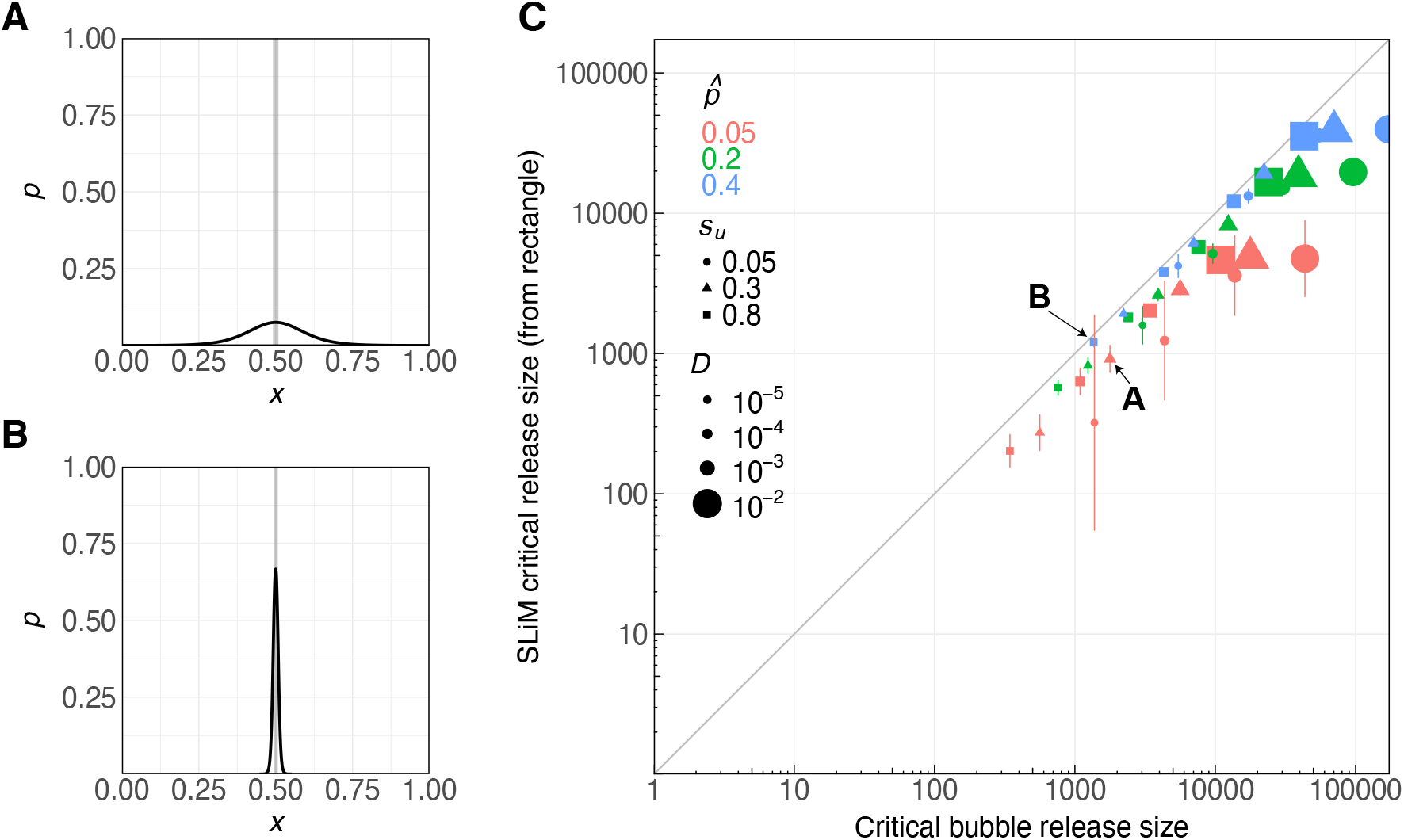
SLiM rectangular release versus reaction-diffusion critical bubble release. **A**. The critical bubble of an underdominance system with *D* = 10^−4^, 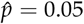, and *s*_*u*_ = 0.3 (black curve), compared to a rectangular release shape (an 100% frequency over a local width) with the same area under the curve. **B**. Same as **A**, but for an underdominance system with *D* = 10^−5^, 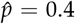, and *s*_*u*_ = 0.8. **C**. Comparison of the critical release size from a rectangular shape in SLiM versus the release size under the critical bubble. Each point represents a different underdominance system, with bars reflecting its transition range in SLiM.

This is not a new finding. Barton and Turelli [43] remarked that introductions of a constant frequency over a certain length could initiate a wave of fixation from a smaller release size than the critical bubble. They referred to such release shapes as “critical initial distributions” and explained that these would always evolve towards the critical bubble. To explore this idea, we used the R package deSolve [53] to simulate the spread of an underdominance system from a rectangular release shape based on its reaction-diffusion equation (Eq. 1; [43]). Figure 4A shows one such wave followed over several time steps. Immediately after the underdominance system is released, the wave falls at the center and spreads out at the sides. In SLiM, this would correspond to individuals with the underdominance allele moving outside of the release width, mating with local wild-types, and passing the allele to offspring. After several time steps, the wave evolves towards the critical bubble but stays slightly above it. In SLiM, several rounds of dispersal, mating, and selection would have increased the number of underdominance alleles during this time period. This explains why critical release sizes under the rectangular shape were consistently less than release sizes under the critical bubble: as the wave moves from the rectangle towards the bubble, the underdominance system accumulates additional alleles, keeping it above the threshold. After remaining near the bubble for many time steps, the wave eventually spreads to fixation.

**Figure 4:**
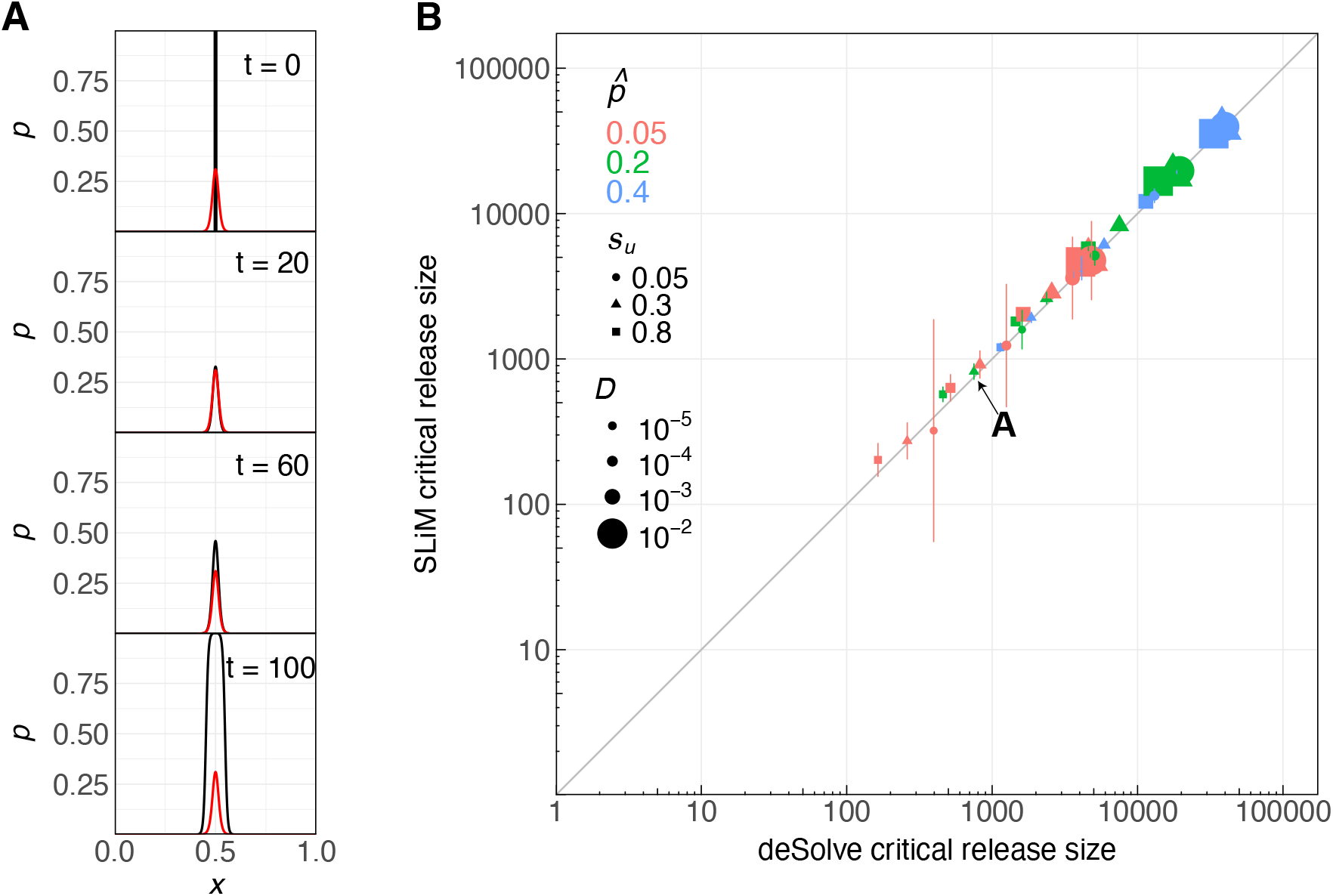
Critical release size from a rectangular release shape. **A**. Evolution of the underdominance allele frequency distribution over time (black curve) from a critical rectangular release at 100% frequency over the critical release width, as predicted by deSolve. The red curve shows the critical bubble distribution for the given underdominance system. **B**. The critical release size in SLiM versus the size predicted by reaction-diffusion simulations in deSolve for a rectangular release of the underdominance allele. Each point represents a different underdominance system, with bars showing its transition range in SLiM.

Using deSolve simulations, we numerically estimated the minimum release width of a rectangular release at which a given underdominance system spreads and defined the deSolve critical release size as the SLiM carrying capacity (100,000) times this width. Comparing this deSolve rectangular critical release size to the SLiM rectangular critical release size, we see much more similar results, with the SLiM transition range encompassing the deSolve prediction for almost all underdominance systems (Figure 4C). In both models, the critical release size was lowest in populations with low *D*. This relates to the pushed-wave dynamics of the underdominance system [43, 44, 54–57]. At the boundaries between underdominance allele and wild-type carriers, the underdominance allele is only present at low frequencies in heterozygotes, who have the lowest fitness. To advance the wave, there must be enough spillover of homozygotes inside the wave into the boundary region. It is easier for these homozygotes to accumulate when underdominance allele carriers remain close together and mate with one another after the release, which occurs more often at low *D*. Critical release sizes also increased with 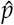 and *s*_*u*_, since systems with higher thresholds and weaker selective advantages require a larger amount of spillover to push underdominance alleles above the threshold at the boundaries.

Our release rectangle optimizes spread by introducing individuals at *p*_0_ = 1 over a sufficiently large region. However, we can apply the same approach to find the critical release width of rectangles with different heights (*p*_0_) or widths. Figure S2 shows the local introduction frequency (*p*_0_) that must be exceeded for an underdominance system to spread from a given release width based on deSolve simulations. In the limiting case where the allele is released everywhere, we obtain the panmictic prediction where *p*_0_ simply must exceed 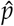. As the release width decreases from one, *p*_0_ must increase. Eventually, the release width becomes so small that the underdominance allele must be released at a 100% local frequency in order to spread. For any release widths below this value, the wave will always contract, assuming density is constant across space. As *D* increases, rectangles must increase in area.

## 6 Homing underdominance gene drives

The critical bubble distribution was derived for an idealized underdominance system with genotype fitness values *f*_*ww*_ = 1, *f*_*dd*_ = 1 + 2*s*_*d*_, and *f*_*dw*_ = 1 + *s*_*d*_ − *s*_*u*_. To achieve a panmictic invasion threshold of 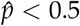—which is typically a prerequisite for spread in a continuous-space population [43]—homozygotes for the underdominance allele must actually be fitter than both wild-type homozygotes and heterozygotes. At present, it remains unclear how such a system could be reliably constructed in practice.

Engineered CRISPR-based gene drive systems, by contrast, may offer a practical means to achieve arbitrarily low panmictic invasion thresholds, even when drive-allele carriers are less fit than wild-type homozygotes, as is likely the case in reality [58]. These systems typically spread through alternative mechanisms, such as homing or toxin–antidote effects. However, their population dynamics are then no longer captured by the reaction term in Equation 1; thus, the critical bubble solution derived in Barton and Turelli [43] no longer applies.

To extend the concept of a critical bubble to a simple threshold-dependent gene drive system, we first consider a single-locus homing underdominance drive with a drive (*d*) and wild-type (*w*) allele. We assume that the drive allele copies itself onto the homologous chromosome in the germline with perfect efficiency (i.e., a cleavage and homing success rate of 100%, with no resistance). Genotype fitness values are parameterized as *f*_*dd*_ = 1 − *s*_*g*_, *f*_*dw*_ = 1 − *hs*_*g*_, and *f*_*ww*_ = 1, where *s*_*g*_ > 0. Unless *s*_*g*_ is very large (e.g., [44, 59]), fitness costs must be greater in heterozygotes than in homozygotes (i.e., *h* > 1) for the drive to display threshold dynamics. Although such a configuration may be challenging to engineer in practice, we use the homing underdominance drive here as a conceptual starting point before exploring more complex architectures.

In Text S2, we derive a reaction–diffusion equation for this system and show that, under the assumption of small drive fitness costs (*s*_*g*_ *≪* 1), an analytical expression for the critical bubble can still be obtained (Equation S4). However, this expression could no longer be integrated to yield a closed-form solution for the area under the critical bubble. We therefore again used deSolve to numerically estimate the critical release size for this drive in a reaction-diffusion model with a rectangular release shape. Figure S3 compares these predictions to SLiM simulation results. For drives with higher invasion thresholds (e.g., 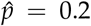), critical release sizes were similar across frameworks, while low-threshold systems (e.g., 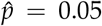) exhibited substantial stochasticity in individual-based simulations, with wide transition ranges; in some replicates, the drive could spread from a release of only 50 individuals (representing a global frequency of just 0.05%). This is again attributable to genetic drift allowing the drive to locally rise above 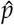, which occurs more readily for homing drives than for underdominance alleles because the former can increase their frequency in the germline. Transition ranges were the widest for drives with minimal fitness costs (e.g., *s*_*g*_ = 0.01), where drift outweighed the effects of weak selection, consistent with previous studies of similar threshold-dependent systems [52, 60–62].

## 7 TADE drive in 2D populations

CRISPR-based toxin-antidote drives offer a powerful and practical approach for modification and suppression drives with threshold-dependent invasion dynamics [28, 63]. Figure 5 provides an illustration of the Toxin-Antidote Dominant Embryo (TADE) drive, a two-locus system involving drive (*d*) and wild-type (*w*) alleles at the drive locus and undisrupted (*a*) and disrupted (*A*) alleles at a target locus on a separate chromosome. The drive construct contains a recoded copy of the target gene that cannot be cut. This target gene is haplolethal, meaning individuals require two functional copies of the gene to survive, either from an undisrupted target allele or the drive allele. Unlike homing drives, TADE does not spread by copying itself onto other chromosomes; instead, it increases its relative frequency by disrupting target alleles and making survival largely dependent on inheriting a drive allele. If drive carriers incur a fitness cost (separate from the toxin effect of the drive), then TADE becomes threshold-dependent [28, 63].

**Figure 5:**
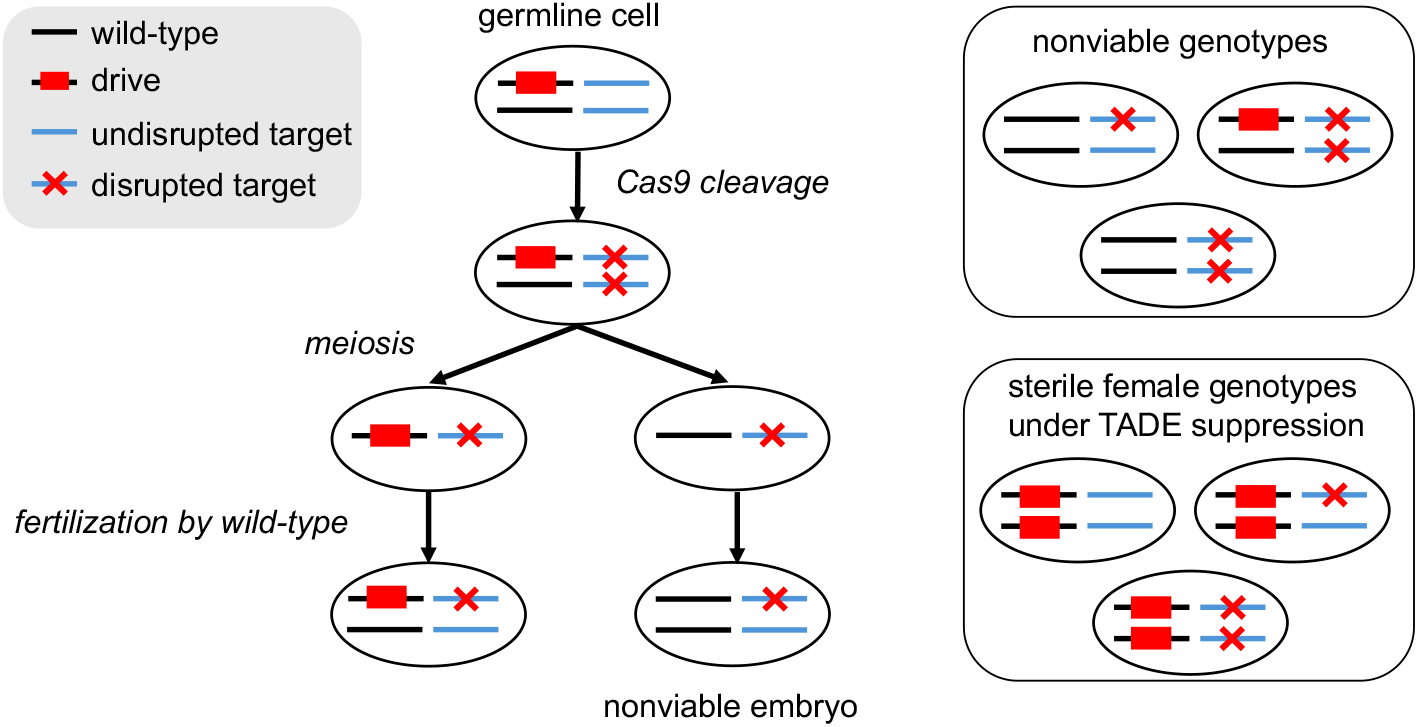
Toxin-Antidote Dominant Embryo (TADE) drive. In the germline of a drive carrier, the drive cleaves and disrupts a haplolethal target gene, but the drive construct also includes a recoded copy of the target gene that is immune to cleavage. Individuals with less than two functional copies of the target gene are nonviable. When TADE is used for population suppression, the drive is inserted into an essential haplosufficient female fertility gene in a manner that disrupts its function, such that females with two drive alleles are sterile.

We implemented a model of the TADE drive assuming that drive carriers cleave undisrupted target alleles at a 100% rate to produce disrupted target alleles, and that cleavage is completely restricted to the germline. For simplicity, we also neglect resistance. Table S2 lists all nine genotypes and their associated fitness cost in our model. Individuals with less than two functional copies of the target gene have fitness 0 and die at the embryo stage. In addition, we assume that there is a fitness cost *s*_*g*_ in drive homozygotes and *s*_*g*_/2 in drive heterozygotes. The panmictic invasion threshold 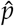 is then a function of *s*_*g*_ and can be found numerically. To model a TADE drive for population modification, we assume that there is a payload gene attached to the drive construct that confers a desirable phenotype. To model a TADE drive for population suppression, we assume that the drive is inserted into a haplosufficient female fertility gene.

Text Text S3 and Tables S2–S5 present detailed derivations of reaction-diffusion equations for the TADE modification and suppression drives. Owing to the complexity of the TADE architecture, a mathematical derivation of the critical bubble—such as that obtained for the homing underdominance drive—is no longer tractable. Nevertheless, we can use numerical reaction–diffusion simulations to estimate the critical release sizes of TADE drives and compare these predictions to the drives’ invasiveness in SLiM models. Moreover, we perform these analyses for the first time in a two-dimensional landscape, taking a further step toward spatial realism.

We extended our SLiM model into 2D by making three modifications to the 1D model: (i) individuals were assigned both *x* and *y* coordinates in a 1×1 square arena; (ii) dispersal distances in the *x* and the *y* directions were drawn independently from a Gaussian with mean 0 and variance *σ*^2^ = 2*D*, yielding an average dispersal distance of 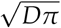; and (iii) the drive was released in a central circle. For each drive, we varied the size of the release circle and determined the minimum diameter that allowed for spread in at least 50% of SLiM replicates. The carrying capacity was held constant at 100,000. Because individuals now inhabit a 1×1 square rather than a 1-unit line in 1D, equilibrium population densities are lower in the 2D model. As before, individuals mate with their nearest neighbor within a circle containing 100 individuals at equilibrium.

Since female drive homozygotes are sterile for the TADE suppression drive, we released drive heterozygotes (*dwAa*) in these simulations, rather than homozygotes as in previous sections. For consistency, heterozygous releases were also used for the TADE modification simulations.

### 7.1 TADE modification

Figure 6 compares the 2D critical release size in SLiM with that predicted by deSolve for the TADE modification drive. For most values of *s*_*g*_ and *D*, SLiM and deSolve predictions were in close agreement. However, for the lowest threshold drive (with *s*_*g*_ = 0.05, such that 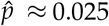), the drive spread readily in SLiM from release circles containing only a handful of drive heterozygotes. This behavior parallels that of the homing underdominance gene drive with 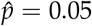 described in Section 6, which effectively behaved as a zero-threshold drive.

**Figure 6:**
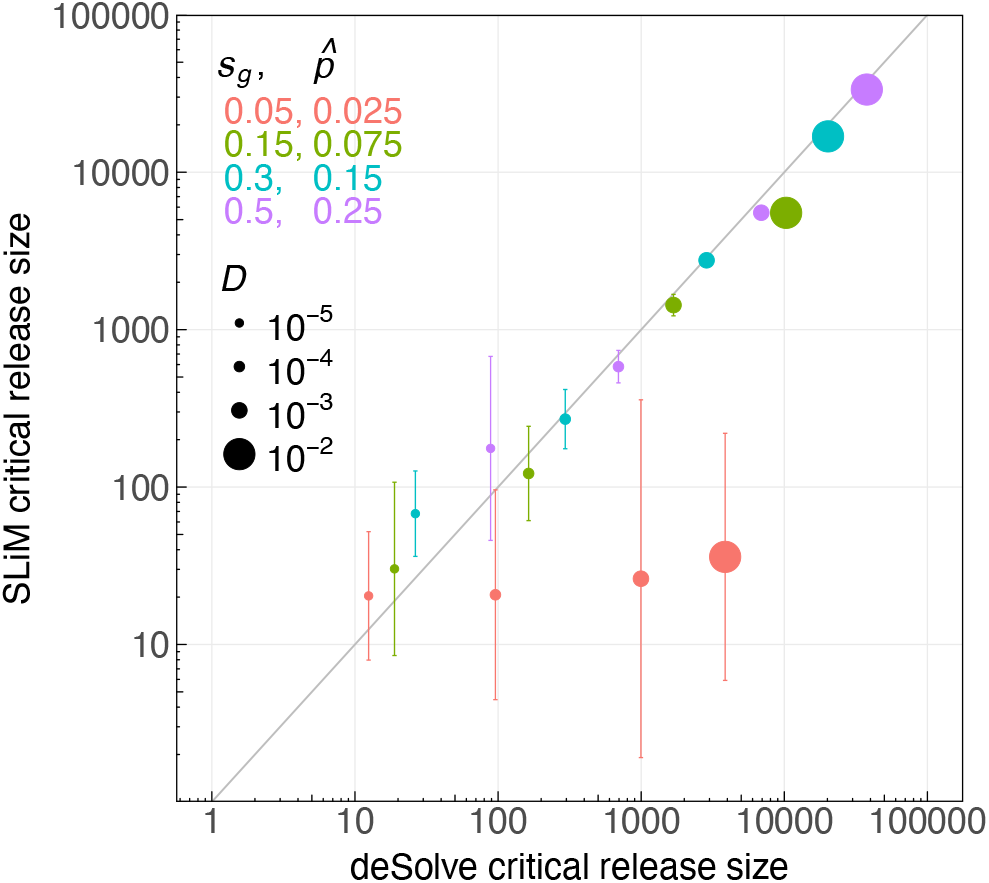
Critical circular releases in SLiM versus deSolve for TADE modification drives. TADE heterozygotes (genotype *dwAa*) were released from a central circle in the 2D SLiM simulations and in 2D deSolve reaction-diffusion model. Each point represents a different TADE modification drive, with bars marking its transition range in SLiM. The panmictic invasion threshold 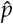 was found numerically, based on the drive’s fitness cost *s*_*g*_ (see Text S3).

For the same values of *D* and *s*_*g*_, TADE modification drives required a larger diameter in 2D than their corresponding width in 1D (Figures S4 and S5). This is because of the increased amount of diffusion in 2D compared to 1D. In 1D, dispersal only occurs along the *x* axis, while in 2D, additional dispersal along the *y* axis results in a larger average dispersal distance 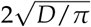 versus 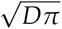, respectively). As a result, the drive wave breaks apart more quickly in 2D. Additionally, since individuals can move in multiple directions, wild-types can invade the release circle more easily, reducing the drive frequency within the wave interior. Lewis and Kareiva analyzed a related problem of determining the critical propagule size required for ecological invasion in populations with an Allee effect [65]. They found that more corrugated (i.e., wavy) boundaries facilitated faster spread than smooth ones. This suggests that release geometry in 2D plays a critical role: threshold-dependent drives could potentially invade from even smaller releases than those shown in Figure 6 if introduced in a shape with a larger perimeter [43, 64].

### 7.2 TADE suppression

We next tested a TADE drive configured for population suppression, in which females with two drive alleles are sterile. Zhang and Champer [63] modeled the spread of such a drive in a 1D reaction-diffusion model and found that it could eliminate a target population but required a relatively large release size to do so. This is because the bulk of drive carriers within the release area quickly self-eliminate due to the absence of fertile females. The drive can then spread only through drive heterozygotes at the boundaries, who suffer from a lower fitness than wild-types. A modification drive, by contrast, can also spread through drive homozygotes pushing outward from the release area. In this section, we extend this analysis to two dimensions and compare the outcomes of deSolve reaction-diffusion simulations with individual-based SLiM simulations.

In our 2D deSolve simulations, heterozygotes mate with each other after the release, creating drive homozygotes in the first generation, who largely self-eliminate—similar to the pattern observed in 1D [63]. This produces an essentially empty release circle in the second generation, with mainly drive heterozygotes along the perimeter in a ring-like shape. However, even a thin ring can still expand in the reaction–diffusion model, as long as enough drive carriers are released for the drive allele to remain above 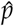 along the perimeter.

In our individual-based SLiM simulations, we observe fundamentally different dynamics, where the drive often fails to eliminate the population under parameter settings for which the reaction–diffusion model predicts success. We can qualitatively distinguish four categories of outcomes in the SLiM simulations: (i) population persistence with complete drive loss, (ii) population persistence with the drive remaining at a low equilibrium frequency, (iii) population persistence with perpetual chasing cycles [47], and (iv) successful population elimination. Which outcome prevails in a given scenario depends primarily on the diffusion constant *D* and the drive fitness cost *s*_*g*_, which determines the panmictic invasion threshold 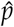.

Figure 7 shows examples of the four different outcomes in our SLiM model for drives that all spread and eliminate the population in the reaction–diffusion model. Complete drive loss tends to occur in scenarios with low *D* and high *s*_*g*_ (Figure 7A). Here, drive carriers do not disperse far from the release circle before homozygotes self-eliminate across the release area, leaving only a thin ring of mostly heterozygotes along the perimeter. Wild-types can then easily penetrate this ring and quickly repopulate the empty area inside it due to reduced competition, whereas the drive is lost. For drives with lower *s*_*g*_—and thus a lower 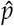 —drive alleles can persist in the population at a low equilibrium frequency even after the release area has been repopulated by wild-type individuals (Figure 7B).

**Figure 7:**
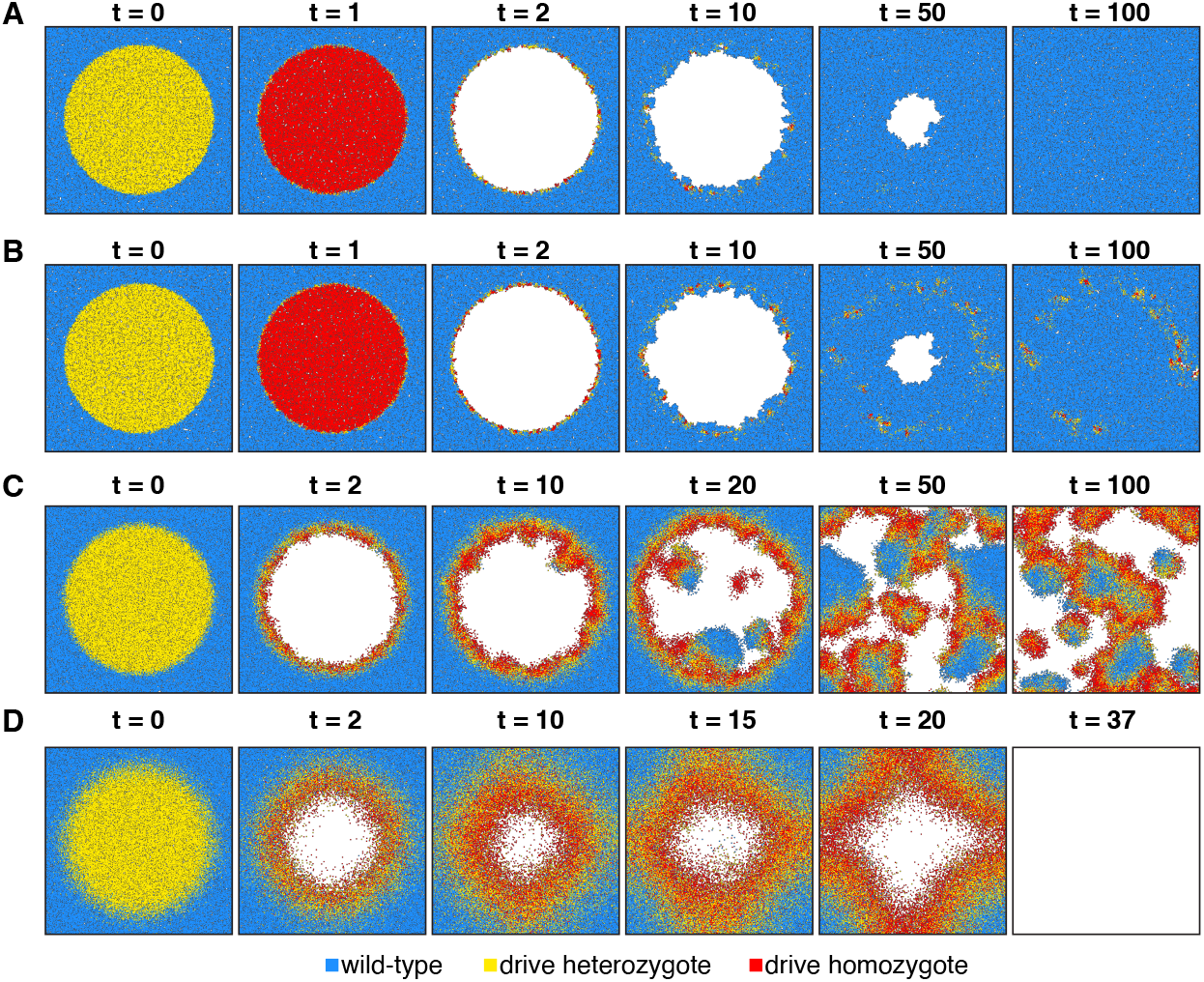
TADE suppression drive outcomes in the 2D SLiM model. Snapshots from four different simulation scenarios in which 50,000 heterozygotes (with genotype *dwAa*) were released in a central circle. Each dot represents an individual and *t* denotes the number of generations since the release. **A**. Simulation with *D* = 10^−5^, *s*_*g*_ = 0.3, and 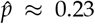. Only a thin ring of drive carriers remains after the initial elimination of most individuals within the release circle. This ring can be easily penetrated by wild-type individuals, and the drive is ultimately lost. **B**. Simulation with *D* = 10^−5^, *s*_*g*_ = 0.05, and 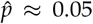. As in **A**, wild-types penetrate the release circle, causing the drive frequency to decline. However, since the drive is only weakly deleterious and more efficient, it persists at a low equilibrium frequency in the population. **C**. Simulation with *D* = 10^−4^, *s*_*g*_ = 0.05, and 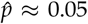. Due to increased dispersal, the ring around the release circle is thicker, with a greater number of drive heterozygotes at the leading edge and drive homozygotes behind. Wild-types can still penetrate the circle and benefit from reduced competition in the emptied central area. The drive “chases” after these new wild-type individuals, emptying these areas once again; however, these dynamics continue perpetually. The population is reduced but not eliminated, yet the drive remains present. **C**. Simulation with *D* = 10^−3^, *s*_*g*_ = 0.3, and 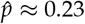. With even higher dispersal, the drive wave becomes thick enough that it can no longer be penetrated by wild-type individuals and expands radially, ultimately eliminating the population.

In scenarios with higher *D* and lower *s*_*g*_ (Figure 7C), the drive wave becomes thicker. Wild-types are still able to penetrate the release circle, but the drive is now efficient enough to spread into these newly rebounding wild-type areas, resulting in chasing dynamics that can continue indefinitely [47, 66]. In these scenarios, the population density tends to remain below carrying capacity but exhibits large fluctuations across time and space. In general, lower-threshold systems reduce population size the most, as such drives can more easily invade new wild-type regions. In contrast, higher-threshold systems spread more slowly and require more drive carriers to eliminate wild-type regions, achieving less population reduction over the same time period. Finally, for even higher values of *D*, drive-carriers can spread so quickly after release that the wave remains wide enough to prevent wild-type individuals from recolonizing the area inside the ring, resulting in complete population elimination (Figure 7D).

## 8 Discussion

In a panmictic population, the invasiveness of a threshold-dependent gene drive system is determined by a single parameter: its invasion threshold frequency 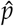. The drive will tend to spread if enough drive alleles are introduced to raise their overall frequency in the population above this threshold. In a continuous-space population with local dispersal, by contrast, the same drive system can spread from a much smaller overall population frequency than 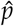, provided that drive alleles are introduced at a sufficiently high local frequency [42–44]. In this study, we examined our ability to predict the conditions under which different threshold-dependent gene drive systems can spread in such populations, a crucial aspect for determining whether a drive can be effectively contained and prevented from invading non-target populations.

We specifically focused on comparing predictions from reaction–diffusion models with those obtained from more realistic individual-based simulations. Diffusion theory offers an elegant mathematical framework in which populations are represented as spatially and temporally varying allele density fields, whose dynamics are governed by differential equations. Yet these models are inherently deterministic, and it has long been recognized that systems composed of discrete individuals can exhibit fundamentally different behaviors [46].

Our results highlight the importance of accounting for the stochasticity in populations composed of discrete individuals when evaluating a gene drive’s invasion potential. For example, while diffusion models predict a sharp threshold separating drive spread from loss, real-world populations invariably exhibit a range of release sizes over which a drive may or may not spread. Reaction–diffusion models can remain valuable for identifying the key parameters that govern invasiveness, and their predicted invasion thresholds often fall within the transition ranges observed in our individual-based simulations. However, their deterministic nature prevents them from capturing the width of these ranges—a limitation that can become critical when the ranges are broad. The most striking discrepancies occurred for suppression drives, which often showed qualitatively different dynamics leading to drive failure under parameter settings for which the diffusion model predicted successful invasion and population elimination.

Mathematical results derived in the reaction-diffusion model—most notably the critical bubble distribution—had previously been obtained only for a simplistic underdominance system in a 1D population [43]. We were able to extend this approach to a simple homing underdominance drive and successfully derived its critical bubble distribution. Unfortunately, the resulting function could no longer be integrated to obtain a closed-form solution for the area under the bubble. This illustrates the mathematical challenges involved in generalizing the critical bubble concept. It remains an open question whether this approach can be successfully extended to other, more realistic types of threshold-dependent drives, such as toxin–antidote systems, which are typically more complex with more loci and more possible genotypes.

Unsurprisingly, we found that threshold-dependent gene drives become more invasive when fitness costs (*s*_*g*_), panmictic invasion thresholds 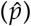, and dispersal rates (*D*) are reduced. However, our results also demonstrate that the spatial configuration of the release plays a critical role. For modification drives that do not substantially reduce population density in the regions they occupy, more spatially concentrated releases generally facilitate spread. This is different for threshold-dependent suppression drives, which can be thwarted by concentrated releases due to what Girardin and Débarre [59] termed “opposing demographic advection.” Here, the drive quickly generates large numbers of homozygotes, which are sterile, causing the advancing wave to collapse (Figure 7). At the wave boundaries, where wild-type individuals greatly outnumber drive carriers, they can readily infiltrate the drive front, leading to loss of the drive or persistent chasing dynamics. These outcomes are consistent with Kläy et al. [66], who showed that once wild-type penetration occurs, the likelihood of chasing increases with dispersal rate and decreases with drive fitness costs. A more diffuse release could, in principle, mitigate this problem by reducing the frequency of matings between drive carriers [67], but this strategy would require the local frequency of the drive to exceed 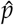 across the release area and for dispersal to be high enough to prevent clustering of drive alleles.

Additional barriers to gene flow could further enhance the containment potential of threshold-dependent drives. Such barriers may be natural (like mountain ranges or water bodies that restrict dispersal) or artificial, such as insecticide-treated zones for mosquito drives [44] or trap arrays for rodent drives. Reaction–diffusion models have shed some light on the critical barrier strength required to prevent a pushed wave from establishing in a new area [44, 62, 68, 69]. These analyses showed that barriers must be strongest for low-threshold systems, which most easily establish, and for populations with high dispersal rates, where dispersal attempts across the boundaries are most frequent. However, stochastic models indicate that if a barrier does not completely block gene flow, genetic drift may occasionally allow alleles to increase in frequency and cross it. Such stochastic breaches are most likely when 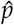 is low and in small populations, where drift is strong [62].

Throughout our analysis, we used simple SLiM models designed to closely follow the assumptions of diffusion theory. As a result, our study did not capture several real-world complexities that could alter drive invasion potential. For instance, we assumed Gaussian dispersal, in which long-distance movement is rare. Frequent long-distance dispersal would likely make it more difficult for a threshold-dependent gene drive to successfully establish, since establishment typically requires many drive carriers to reach the same location simultaneously.

We also modeled a homogeneous environment with uniform carrying capacities and introduced the drive at the same density as the surrounding population. If the drive were released at a higher local density without increasing the local carrying capacity, drive carriers would experience stronger competition in the release area, causing many drive alleles to be quickly eliminated; therefore, we would not expect substantially different results. In contrast, a release in a region of higher carrying capacity could create a positive density gradient that favors drive spread, as drive carriers at the wave front would outnumber wild-types. Indeed, Champer et al. [42] found that elevated carrying capacities in the release region made it easier for drives to spread and sometimes allowed even drives with 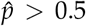 to persist. These findings underscore the importance of detailed modeling of both the target population and potential spillover populations (e.g., ship harbors for rodent suppression drives). Measures that reduce the carrying capacity of such spillover regions could diminish the risk of unintended invasion.

To better understand the invasion criteria of a threshold-dependent gene drive, experimentalists should at a minimum estimate 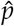 by varying the initial release frequency in cage experiments and identifying the minimum frequency at which the drive consistently spreads. Accurate estimates of drive fitness costs (*s*_*g*_) will also be essential, while acknowledging that they may differ markedly under natural conditions. Combined with life-history traits such as fecundity and lifespan, and ecological parameters such as dispersal rates and population density, these measurements can then inform simulation models that systematically evaluate invasion potential.

Ideally, invasiveness would be studied directly in large, fully contained field trials by releasing the drive at high local frequency and varying the release area. This would reveal the minimum area required for spread and the range over which the probability of spread exceeds a given threshold. Smaller introductions than this threshold would pose little invasion risk, whereas releases over larger areas could enable spread even from lower local frequencies, provided 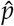 is exceeded within the release region. Such experiments will unfortunately remain impractical for most target species, especially those with high dispersal rates.

Overall, our study highlights the need for realistic spatial modeling when evaluating the risks and containment potential of gene drives. Diffusion theory remains invaluable for identifying key parameters, such as the critical frequency distribution required for spread [43]. Yet even in large individual-based models, stochasticity produced divergent outcomes: instead of a sharp boundary between spread and loss, we observed a broad transition range of release sizes where invasion was probabilistic. These ranges were typically wider for low-threshold, mildly deleterious drives and would likely expand in smaller populations where stochastic effects are stronger. Previous studies have likewise shown that under strong drift, low-threshold systems can behave like zero-threshold systems [52, 61, 62]. For suppression drives, stochasticity can fundamentally alter qualitative outcomes, generating dynamic behaviors such as chasing that are absent from diffusion models. Together, these results demonstrate that both stochasticity and spatial structure are essential considerations for accurately assessing the invasiveness and containment of threshold-dependent gene drives.

## Acknowledgments

The authors thank Florence Débarre for helpful discussions about reaction-diffusion simulations. This work was supported by the National Institutes of Health under award R35GM152242.

## Data and Code Availability

Simulation models and supporting code are available on GitHub (https://github.com/MesserLab/Containment)

## Notes

### Competing Interest Statement

The authors have declared no competing interest.

