## Supplemental Information for "Predicting the invasiveness of threshold-dependent gene drives"

Isabel K. Kim

Philipp W. Messer*

Department of Computational Biology, Cornell University, Ithaca, NY 14853, United States

### Supplemental Text

#### Text S1 Critical bubble for the underdominance system

In this section, we recapitulate the derivation of the critical bubble of an underdominance allele from Barton and Turelli [1].

The underdominance system is characterized by $s_u$, the strength of underdominance selection, and $\hat{p}$, the invasion threshold. The change in allele frequency following one generation of random mating can be described by

$$\Delta p = \frac{s_u p(1-p)(2p+\alpha-1)}{1+2ps_u(\alpha-1+p)}.$$

To derive the critical bubble, one must assume that $s_u$ is small, such that the denominator is approximately 1. This is often referred to as the "cubic approximation" [1], since the reaction equation becomes cubic in $p$, with $\Delta p \approx s_u p(1-p)(2p+\alpha-1)$. Solving this in terms of $\hat{p}$ (with $\alpha = 1-2\hat{p}$) yields $\Delta p \approx 2s_u p(1-p)(p-\hat{p})$.

We define $D$ as the diffusion constant, equal to half the variance in dispersal distances between mother and offspring. In 1D space, the spread of the underdominance system can be defined by the exact reaction-diffusion equation of

$$\begin{aligned}\frac{\partial p(x,t)}{\partial t} &= D\frac{\partial^2}{\partial x^2}p + \frac{s_u p(1-p)(2p+\alpha-1)}{1+2ps_u(\alpha-1+p)}\\ &= D\frac{\partial^2}{\partial x^2}p + \frac{2s_u p(1-p)(p-\hat{p})}{1+2ps_u(p-2\hat{p})},\end{aligned}$$

or the cubic approximation of

$$\frac{\partial p(x,t)}{\partial t} = D\frac{\partial^2}{\partial x^2}p + s_u p(1-p)(2p+\alpha-1). \tag{S1}$$

This equation can similarly be written in terms of $\hat{p}$ (see Eq. 1).

To derive the critical bubble, the authors first rescaled time and space with

$$T = s_u t \quad \text{and}$$

$$X^2 = \frac{s_u}{D}x^2,$$

such that

$$\frac{\partial p}{\partial t} = s_u \frac{\partial p}{\partial T} \quad \text{and}$$
$$\frac{\partial^2 p}{\partial x^2} = \frac{s_u}{D} \frac{\partial^2 p}{\partial X^2}.$$

Plugging these terms into Eq. S1 and canceling out $s_u$ results in a rescaled reaction-diffusion equation of

$$\frac{\partial p}{\partial T} = \frac{\partial^2 p}{\partial X^2} + p(1-p)(2p+\alpha-1).$$

In the critical bubble distribution, the effects of the reaction and diffusion are exactly balanced; therefore, $\partial p/\partial T = 0$. Rearranging this gives

$$0 = \frac{\partial^2 p}{\partial X^2} + p(1-p)(2p+\alpha-1),$$
$$\frac{\partial^2 p}{\partial X^2} = -p(1-p)(2p+\alpha-1).$$

We can solve for $\partial p/\partial X$ by letting $y = \partial p/\partial X$, such that $\partial^2 p/\partial X^2 = y\partial y/\partial p$.

$$y\frac{dy}{dp} = -\,p(1-p)(2p+\alpha-1),$$
$$\int y\,dy = \int -\,p(1-p)(2p+\alpha-1)\,dp,$$
$$\frac{1}{2}y^2 = \frac{1}{2}p^2\left[(1-p)^2 - \alpha\left(1-\frac{2p}{3}\right)\right],$$
$$\frac{\partial p}{\partial X} = p\sqrt{(1-p)^2 - \alpha\left(1-\frac{2p}{3}\right)} \quad \text{(with } p \geq 0\text{)}.$$

Since the critical bubble is unimodal and symmetric, at the center of the bubble ($X = 0$), $\partial p/\partial X = 0$. Solving $\partial p/\partial X$ for the frequency at the center of the curve (represented here with $p_c$) and rejecting frequencies outside of $[0, 1]$ gives

$$p_c(\alpha) = 1 - \frac{\alpha}{3} - \frac{1}{3}\sqrt{\alpha(3+\alpha)},$$

which is represented in terms of $\hat{p}$ in Eq. 2.

Next, we consider the reciprocal of $\partial p/\partial X$:

$$\frac{\partial X}{\partial p} = \frac{1}{p\sqrt{(1-p)^2 - \alpha(1-\frac{2p}{3})}}.$$

Centering the bubble at $X = 0$, the value of $X > 0$ associated with each frequency in the bubble can be found by integrating $\partial X/\partial p$ with respect to $p$, from 0 to $p_c$:

$$X(p) = \frac{1}{\sqrt{1-\alpha}}\log\left\{\frac{3}{p\sqrt{\alpha(3+\alpha)}}\left[1 - p - \alpha(1-\frac{p}{3}) + \sqrt{1-\alpha}\sqrt{(1-p)^2 - \alpha(1-\frac{2p}{3})}\right]\right\}.$$

The critical bubble in *unscaled* space is then found by multiplying $X(p)$ by $\sqrt{D/s_u}$:

$$x(p) = \sqrt{\frac{D}{s_u}}\frac{1}{\sqrt{1-\alpha}}\log\left\{\frac{3}{p\sqrt{\alpha(3+\alpha)}}\left[1 - p - \alpha(1-\frac{p}{3}) + \sqrt{1-\alpha}\sqrt{(1-p)^2 - \alpha(1-\frac{2p}{3})}\right]\right\}. \tag{S2}$$

When this derivation is done in Mathematica [2], an expression that involves an inverse tangent and several $\sqrt{-1+\alpha}$ terms is returned. Since $0 < \alpha < 1$, $\sqrt{-1+\alpha}$ is imaginary; rewriting this as $\sqrt{-1 \times (1-\alpha)} = i\sqrt{1-\alpha}$ and then applying the identity $\tan^{-1}(iz) = i\tanh^{-1}(z)$ converts the expression to exponential form. This yields Eq. S2.

Finally, one can find the area under the critical bubble—proportional to the number of individuals that would need to be released in this distribution—by integrating the scaled-space critical bubble equation with respect to $p$, from 0 to $p_c$:

$$M(\alpha) = \log\left(\frac{\sqrt{\alpha(\alpha+3)}}{3 - \alpha - 3\sqrt{1-\alpha}}\right).$$

Eq. 4 shows $M(\alpha)$ in terms of $\hat{p}$.

Transforming this back to unscaled space gives the area under the curve formula of Eq. 3, which includes a factor of 2 to account for both the right-hand side of the critical bubble ($x > 0$) and left-hand side of the critical bubble ($x < 0$).

#### Text S2 Critical bubble for the homing underdominance gene drive

To derive the critical bubble analytically for a gene drive system, the drive must satisfy several conditions. First, since the critical bubble is a function of only one frequency, the gene drive cannot involve more than two alleles. This eliminates toxin-antidote gene drives (which involve drive alleles, wild-type alleles, and disrupted alleles), two-locus systems, and gene drives with resistance. Second, to use the cubic approximation necessary in the critical bubble derivation (Section Text S2), the fitness cost of the drive (represented here with $s_g$) must be small. Third, the gene drive must be threshold-dependent. For a biallelic gene drive system with low $s_g$, a threshold can only be achieved if heterozygotes have the lowest fitness. Lastly, there must be some mechanism favoring the drive allele such that, when the drive is released above the invasion threshold ($\hat{p}$), the population is pushed towards homozygosity for the drive allele rather than wild-type allele. This can be satisfied through homology-directed repair, which converts wild-type alleles to drive alleles in the germline of drive carriers.

Thus, the critical bubble can only be derived analytically if one assumes a rather unrealistic gene drive system: a homing gene drive with no resistance, a small fitness cost in homozygotes, and the highest fitness cost in heterozygotes. To derive critical bubbles numerically, however, some of these assumptions can be dropped (e.g., Tanaka et al. [3]).

In this section, we use Barton and Turelli [1]'s framework to derive the critical bubble of a homing underdominance gene drive system. We parameterize fitness values with $f_{ww} = 1$, $f_{dd} = 1 - s_g$, and $f_{dw} = 1 - hs_g$ (with $h > 1$) and a germline conversion rate $c$.

We let $p(t)$ represent the frequency of the drive in the germline at generation $t$. After a fraction $c$ of wild-type alleles have been converted to drive alleles in heterozygotes, this frequency becomes $p(t) = p_{dd} + 0.5(1 + c)p_{dw}$.

After random mating, genotype frequencies will be in Hardy-Weinberg equilibrium with respect to $p(t)$. We assume that fitness affects viability selection, so genotype frequencies in the next generation are

$$p_{dd}(t+1) = \frac{p^2(1-s_g)}{\bar{w}},$$
$$p_{dw}(t+1) = \frac{2p(1-p)(1-hs_g)}{\bar{w}}, \quad \text{and}$$
$$p_{ww}(t+1) = \frac{(1-p)^2}{\bar{w}}, \quad \text{with}$$
$$\bar{w}(t+1) = p^2(1-s_g) + 2p(1-p)(1-hs_g) + (1-p)^2 \quad \text{representing the mean fitness.}$$

Germline conversion then occurs in adult heterozygotes, such that the drive frequency in the germline becomes

$$p(t+1) = \frac{p^2(1-s_g)}{\bar{w}} + \frac{p(1-p)(1-hs_g)(1+c)}{\bar{w}}.$$

This results in the following reaction equation:

$$\Delta p = \frac{p(p-1)\left[s_g(h+p-2hp) + c(hs_g-1)\right]}{1+2phs_g(p-1) - s_g p^2}.$$

By setting the reaction equation equal to 0, we can calculate the invasion threshold of the drive ($\hat{p}$) as

$$\hat{p} = \frac{c - hs_g(1+c)}{s_g(1-2h)}. \tag{S3}$$

Thus, the reaction-diffusion equation of the system is

$$\frac{\partial p}{\partial t} = D\frac{\partial^2 p}{\partial x^2} + \frac{p(p-1)\left[s_g(h+p-2hp) + c(hs_g-1)\right]}{1+2phs_g(p-1) - s_g p^2}.$$

To derive the critical bubble, we use the cubic approximation (assuming small $s_g$) and obtain an approximate reaction-diffusion equation of

$$\frac{\partial p}{\partial t} = D\frac{\partial^2 p}{\partial x^2} + p(p-1)\left[s_g(h+p-2hp) + c(hs_g-1)\right].$$

Following the procedure detailed in Text S1, we can use this reaction-diffusion equation to derive the frequency at the center of the critical bubble:

$$\tilde{p} = \frac{\frac{2}{3}(3h + ch - \frac{c}{s_g} - 1) - \sqrt{\frac{4}{9}(\frac{c}{s_g} - ch - 3h + 1)^2 - 2(2h-1)(h + ch - \frac{c}{s_g})}}{2h-1}$$

Finally, we can derive the critical bubble $x(p)$, the unique 1D frequency distribution at which diffusion and selection exactly balance:

$$x(p) = \sqrt{\frac{D}{hs_g + c(hs_g - 1)}} \left\{ \log(1-\theta) - \log(1+\theta) + \log(1+\beta) - \log(1-\beta) \right\}, \tag{S4}$$

with

$$\begin{aligned}\theta = & \frac{1}{3\sqrt{2h-1}\sqrt{s_g}\sqrt{hs_g + c(hs_g-1)}} [s_g\sqrt{2} - 3hs_g\sqrt{2} - c\sqrt{2}(hs_g - 1) \\ & + s_g\sqrt{2 - 3h - \frac{c(6h-5)(hs_g-1)}{s_g} + \frac{2c^2(hs_g-1)^2}{s_g^2}}] \qquad \text{and} \\ \beta = & \frac{\sqrt{s_g}}{3\sqrt{2}\sqrt{chs_g + hs_g - c}} [-3p\sqrt{2h-1} \\ & + \sqrt{18h(p-1)^2 + 3p(4-3p) - \frac{6c(2p-3)(hs_g-1)}{s_g}}.\end{aligned}$$

The area under the curve cannot be easily obtained through mathematical integration but can be found numerically. When the population is distributed uniformly, the total release size under the critical bubble is then just the population size multiplied by the area under the curve.

#### Text S3 Reaction-diffusion model for the TADE drive

In this section, we derive the six reaction-diffusion equations of the Toxin-Antidote Dominant Embryo (TADE) drive (one for each viable genotype frequency). Table S3 provides a description of all symbols used in this section.

TADE involves two loci on separate chromosomes. At the drive locus, two alleles are possible: drive ($d$) and wild-type ($w$). At the target locus, two alleles are also possible: undisrupted ($a$) and disrupted ($A$). This yields nine potential genotypes. We assign a number ($i$) to each genotype in Table S2.

The drive targets a haplolethal gene, such that individuals need two functional copies of the gene to survive. We assume that an undisrupted target site is cleaved at rate $c$ in drive carriers and all cleavage events disrupt the target gene (i.e., there is no resistance). The drive construct contains a functional copy of the target gene that is recoded to be immune to cleavage. Thus, individuals with two drive alleles, two undisrupted target alleles, or one of each are viable, while individuals lacking two functional copies are nonviable with fitness ($f_i$) of 0. This includes $wwAa$ ($i = 4$), $wwAA$ ($i = 7$), and $dwAA$ ($i = 8$).

For the suppression version of the drive, we assume that the drive allele is placed in an essential haplosufficient female fertility gene in a manner that disrupts its function, such that females with two drive alleles are sterile. This includes $ddaa$ ($i = 3$), $ddAa$ ($i = 6$), and $ddAA$ ($i = 9$), resulting in only three viable genotypes that are fertile as females and six viable genotypes that are fertile as males. For the modification version of the drive, all six viable genotypes are assumed to be fertile in both sexes.

For TADE modification, we track the frequencies of each genotype ($p_i$), assuming constant density across space. In contrast, the TADE suppression drive is designed to eliminate the population and therefore strongly impacts population density. Thus, for TADE suppression, we track genotype densities ($N_i$) instead.

Table S4 lists all 21 possible mating crosses between viable genotypes, along with their probabilities for TADE modification and TADE suppression, assuming random mating and an equal sex ratio. We denote the probability of cross $m$ with $\gamma_m$.

Under the TADE modification drive, all genotypes are fertile, so a cross between genotype $a$ and $b$ (with $a \neq b$) is possible if genotype $a$ is male and genotype $b$ is female or vice versa, resulting in a factor 2 in these $\gamma_m$. However, under the TADE suppression drive, genotypes $i = 3$, 6, and 9 are sterile as females, so mating crosses involving these genotypes are only possible if these individuals are male and the other individual is a fertile female. This results in only a factor 1 in $\gamma_m$ of such crosses and $\gamma_m = 0$ for crosses involving two of genotype $i = 3$, 6, or 9. The denominator of $\gamma_m$ for TADE suppression accounts for the total density of fertile females and the total density of fertile males.

We next consider the probability of each offspring genotype $i$ resulting from mating cross $m$, denoting this with $\omega_i(m)$. Table S5 lists all $\omega_i(m)$ for both TADE modification and suppression.

We assume that adults are removed after random mating, creating discrete generations. Offspring then experience viability selection, with survival probabilities equal to their genotype-based fitness ($f_i$).

#### *Text S3.1 TADE modification*

For TADE modification, we derive the frequency of genotype $i$ in the next generation ($t+1$) by summing over the probability of each possible mating cross at time $t$, weighted by the probability that an offspring of such a cross has genotype $i$. We then consider the fitness of genotype $i$ and normalize by the mean fitness of all offspring, $\bar{f}(t+1)$:

$$\bar{f}(t+1) = \sum_{j=1}^{9} \left[ f_j \sum_{m=1}^{21} \gamma_m(t) \omega_j(m) \right], \tag{S5}$$

$$p_i(t+1) = \frac{f_i \sum_{m=1}^{21} \gamma_m(t) \omega_i(m)}{\bar{f}(t+1)}.$$

This results in a reaction equation for genotype $i$ of

$$f(p_i) = p_i(t+1) - p_i(t).$$

To simulate changes in genotype frequency across continuous time and space, we add a diffusion term. In 1D, the reaction-diffusion equation becomes

$$\frac{\partial p_i}{\partial t} = D \frac{\partial^2 p_i}{\partial x^2} + f(p_i).$$

And in 2D, the reaction-diffusion equation is

$$\frac{\partial p_i}{\partial t} = D \frac{\partial^2 p_i}{\partial x^2} + D \frac{\partial^2 p_i}{\partial y^2} + f(p_i).$$

The drive allele frequency ($p_d(t)$) is calculated using the frequencies of the five viable genotypes with the drive allele:

$$p_d(t) = \frac{1}{2}[p_2(t) + p_5(t)] + p_3(t) + p_6(t) + p_9(t)$$

The reaction equation for the drive allele is then:

$$f(p_d) = p_d(t+1) - p_d(t).$$

The invasion threshold ($\hat{p}$; where $f(p_d) = 0$) does not have a closed-form expression but can be found numerically, given the genotypes introduced.

The reaction-diffusion equation for the drive allele is defined following the same approach as before. In 1D:

$$\frac{\partial p_d}{\partial t} = D\frac{\partial^2 p_d}{\partial x^2} + f(p_d).$$

And in 2D:

$$\frac{\partial p_d}{\partial t} = D\frac{\partial^2 p_d}{\partial x^2} + D\frac{\partial^2 p_d}{\partial y^2} + f(p_d).$$

#### *Text S3.2 TADE suppression*

For the TADE suppression drive, we focus on genotype densities ($N_i$) rather than frequencies ($p_i$). $N_i$ is affected not only by reproduction and viability selection but also by density-dependent competition. We assume that competition for resources occurs within a radius of $dx$. In an arena with area 1 and capacity 100,000, the local carrying capacity is equal to $\kappa = 100000 \times dx$. Individuals are more likely to survive competition when there are fewer individuals in the region and less likely when there are more. If the total number of individuals before competition at time $t+1$ is $N'_{tot}(t+1)$, the probability of surviving competition ($\psi(t+1)$) is

$$\psi(t+1) = \min(\frac{\kappa}{N'_{tot}(t+1)}, 1).$$

If the population density is below the local carrying capacity, all individuals survive, and if the population density exceeds the local carrying capacity, only $\kappa$ individuals survive.

To calculate $N'_{tot}$, we assume that each fertile female produces $L$ offspring and the overall probability of offspring surviving viability selection is equal to the mean fitness of offspring, $\bar{f}(t+1)$, as given by Eq. S5:

$$N'_{tot}(t+1) = \frac{1}{2}(N_1(t) + N_2(t) + N_5(t))(L)(\bar{f}(t+1)).$$

After competition, the total population density is

$$N_{tot}(t+1) = \psi(t+1) \times N'_{tot}(t+1).$$

We calculate the density of genotype $i$ in the next generation using the same approach as in the previous section for the TADE modification drive: we sum over the probability of each mating cross in the previous generation ($\gamma_m(t)$) and the probability that offspring of each cross had genotype $i$ ($\omega_i(m)$) and then account for the probability of such offspring surviving viability selection ($f_i$):

$$N_i(t+1) = \left( \frac{f_i \sum_{m=1}^{21} \gamma_m(t)\omega_i(m)}{\bar{f}(t+1)} \right) \left( N_{tot}(t+1) \right).$$

The reaction equation for genotype $i$ is then

$$f(N_i) = N_i(t+1) - N_i(t).$$

Adding a diffusion term to model the spread of genotype $i$ across continuous space and time gives, in 1D:

$$\frac{\partial N_i}{\partial t} = D\frac{\partial^2 N_i}{\partial x^2} + f(N_i)$$

and in 2D:

$$\frac{\partial N_i}{\partial t} = D\frac{\partial^2 N_i}{\partial x^2} + D\frac{\partial^2 N_i}{\partial y^2} + f(N_i).$$

To calculate the total density of drive alleles at time $t$, we sum over the densities of each genotype, weighted by the number of drive alleles each genotype possesses:

$$N_d(t) = N_2(t) + N_5(t) + 2[N_3(t) + N_6(t) + N_9(t)]. \tag{S6}$$

The frequency of the drive allele at $t$ can then be found by dividing the drive allele density by the total allelic density:

$$p_d(t) = \frac{N_d(t)}{2[N_1(t) + N_2(t) + N_3(t) + N_5(t) + N_6(t) + N_9(t)]}$$

The invasion threshold ($\hat{p}$; where $p_d(t+1) - p_d(t) = 0$) again cannot be expressed analytically but can be determined numerically, given the genotypes introduced.

The reaction-diffusion equation for the drive allele density is defined based on the current densities of all genotypes (Eq. S6). In 1D:

$$\frac{\partial N_d}{\partial t} = D\frac{\partial^2 N_d}{\partial x^2} + f(N_d).$$

And in 2D:

$$\frac{\partial N_d}{\partial t} = D\frac{\partial^2 N_d}{\partial x^2} + D\frac{\partial^2 N_d}{\partial y^2} + f(N_d).$$

### Supplemental Tables

**Table S1: Summary of variables used throughout the study.** Not all variables apply to every system modeled. For systems other than the underdominance allele, drive cleavage is assumed to occur in the germline and is followed by drive conversion for the homing underdominance gene drive or target gene disruption for the TADE drive, with no resistance.

| Variable | Description | Note |
|---|---|---|
| **Shared variables** | | |
| $D$ | Diffusion constant | Half the variance in displacement along the $x$ or $y$ axis |
| $\hat{p}$ | Panmictic invasion threshold | |
| $p_0$ | Local introduction frequency | Within the release shape |
| **Underdominance allele** | | |
| $s_u$ | Underdominance selection coefficient | |
| $s_d$ | Directional selection coefficient | |
| $\alpha$ | Ratio of directional to underdominance selection | $\hat{p} = (1-\alpha)/2$ |
| $\phi$ | Multiplicative factor applied to critical bubble | See Section 4 |
| **Homing underdominance gene drive & TADE drive** | | |
| $s_g$ | Fitness cost of the drive allele | |
| $h$ | Dominance coefficient of the drive allele | $h > 1$ for homing underdominance, $h = 1/2$ for TADE |
| $c$ | Drive cleavage rate | $c = 1$ |

**Table S2: Representation and fitness of TADE genotypes.** † indicates genotypes that are sterile as females under the TADE suppression drive.

| Index ($i$) | Genotype | Fitness ($f_i$) |
|---|---|---|
| 1 | $wwaa$ | 1 |
| 2 | $dwaa$ | $1 - hs_g$ |
| 3 | $ddaa^\dagger$ | $1 - s_g$ |
| 4 | $wwAa$ | 0 |
| 5 | $dwAa$ | $1 - hs_g$ |
| 6 | $ddAa^\dagger$ | $1 - s_g$ |
| 7 | $wwAA$ | 0 |
| 8 | $dwAA$ | 0 |
| 9 | $ddAA^\dagger$ | $1 - s_g$ |

**Table S3: Additional variables used in TADE derivations.**

| Variable | Description |
|---|---|
| $i$ | Genotype index |
| $p_i$ | Frequency of genotype $i$ |
| $p_d$ | Drive allele frequency |
| $N_i$ | Density of genotype $i$ |
| $N_d$ | Total density of drive alleles |
| $f_i$ | Fitness of genotype $i$ |
| $\bar{f}$ | Population mean fitness |
| $m$ | Mating cross index |
| $\gamma_m$ | Probability of mating cross $m$ |
| $\omega_i(m)$ | Probability of offspring of mating cross $m$ having genotype $i$ |
| $dx$ | Local area over which competition occurs |
| $\kappa$ | Local carrying capacity |
| $N'_{tot}$ | Total density of individuals before density-dependent competition |
| $\psi$ | Probability of surviving competition |
| $L$ | Litter size |
| $N_{tot}$ | Total density of individuals after density-dependent competition |

**Table S4: Probability of TADE mating crosses.** $m$ is the index of the mating cross, and $\gamma_m$ is the probability of the cross. Probabilities are shown separately for TADE modification drives and TADE suppression drives.

| Cross ($m$) | $\gamma_m$ for TADE modification | $\gamma_m$ for TADE suppression |
|---|---|---|
| $wwaa \times wwaa$ | $p_1^2$ | $\frac{N_1^2}{(N_1+N_2+N_5)(N_1+N_2+N_3+N_5+N_6+N_9)}$ |
| $wwaa \times dwaa$ | $2p_1p_2$ | $\frac{2N_1N_2}{(N_1+N_2+N_5)(N_1+N_2+N_3+N_5+N_6+N_9)}$ |
| $wwaa \times ddaa$ | $2p_1p_3$ | $\frac{N_1N_3}{(N_1+N_2+N_5)(N_1+N_2+N_3+N_5+N_6+N_9)}$ |
| $wwaa \times dwAa$ | $2p_1p_5$ | $\frac{2N_1N_5}{(N_1+N_2+N_5)(N_1+N_2+N_3+N_5+N_6+N_9)}$ |
| $wwaa \times ddAa$ | $2p_1p_6$ | $\frac{N_1N_6}{(N_1+N_2+N_5)(N_1+N_2+N_3+N_5+N_6+N_9)}$ |
| $wwaa \times ddAA$ | $2p_1p_9$ | $\frac{N_1N_9}{(N_1+N_2+N_5)(N_1+N_2+N_3+N_5+N_6+N_9)}$ |
| $dwaa \times dwaa$ | $p_2^2$ | $\frac{N_2^2}{(N_1+N_2+N_5)(N_1+N_2+N_3+N_5+N_6+N_9)}$ |
| $dwaa \times ddaa$ | $2p_2p_3$ | $\frac{N_2N_3}{(N_1+N_2+N_5)(N_1+N_2+N_3+N_5+N_6+N_9)}$ |
| $dwaa \times dwAa$ | $2p_2p_5$ | $\frac{2N_2N_5}{(N_1+N_2+N_5)(N_1+N_2+N_3+N_5+N_6+N_9)}$ |
| $dwaa \times ddAa$ | $2p_2p_6$ | $\frac{N_2N_6}{(N_1+N_2+N_5)(N_1+N_2+N_3+N_5+N_6+N_9)}$ |
| $dwaa \times ddAA$ | $2p_2p_9$ | $\frac{N_2N_9}{(N_1+N_2+N_5)(N_1+N_2+N_3+N_5+N_6+N_9)}$ |
| $ddaa \times ddaa$ | $p_3^2$ | 0 |
| $ddaa \times dwAa$ | $2p_3p_5$ | $\frac{N_3N_5}{(N_1+N_2+N_5)(N_1+N_2+N_3+N_5+N_6+N_9)}$ |
| $ddaa \times ddAa$ | $2p_3p_6$ | 0 |
| $ddaa \times ddAA$ | $2p_3p_9$ | 0 |
| $dwAa \times dwAa$ | $p_5^2$ | $\frac{N_5^2}{(N_1+N_2+N_5)(N_1+N_2+N_3+N_5+N_6+N_9)}$ |
| $dwAa \times ddAa$ | $2p_5p_6$ | $\frac{N_5N_6}{(N_1+N_2+N_5)(N_1+N_2+N_3+N_5+N_6+N_9)}$ |
| $dwAa \times ddAA$ | $2p_5p_9$ | $\frac{N_5N_9}{(N_1+N_2+N_5)(N_1+N_2+N_3+N_5+N_6+N_9)}$ |
| $ddAa \times ddAa$ | $p_6^2$ | 0 |
| $ddAa \times ddAA$ | $2p_6p_9$ | 0 |
| $ddAA \times ddAA$ | $p_9^2$ | 0 |

**Table S5: Offspring genotype probabilities for each TADE mating crosses.** Each row corresponds to a mating cross $m$ between two parental genotypes, and each column gives the probability that the offspring has genotype $i$. The † denotes crosses that are impossible (i.e., $\gamma_m = 0$) under the TADE suppression drive. See Table S4 for the probabilities of each cross ($\gamma_m$) and Table S3 for genotype indices ($i$).

| Cross ($m$) | $\omega_1(m)$ | $\omega_2(m)$ | $\omega_3(m)$ | $\omega_4(m)$ | $\omega_5(m)$ | $\omega_6(m)$ | $\omega_7(m)$ | $\omega_8(m)$ | $\omega_9(m)$ |
|---|---|---|---|---|---|---|---|---|---|
| $wwaa \times wwaa$ | 1 | 0 | 0 | 0 | 0 | 0 | 0 | 0 | 0 |
| $wwaa \times dwaa$ | $\frac{1}{2}(1-c)$ | $\frac{1}{2}(1-c)$ | 0 | $\frac{1}{2}c$ | $\frac{1}{2}c$ | 0 | 0 | 0 | 0 |
| $wwaa \times ddaa$ | 0 | $1-c$ | 0 | 0 | $c$ | 0 | 0 | 0 | 0 |
| $wwaa \times dwAa$ | $\frac{1}{4}(1-c)$ | $\frac{1}{4}(1-c)$ | 0 | $\frac{1}{4}(1+c)$ | $\frac{1}{4}(1+c)$ | 0 | 0 | 0 | 0 |
| $wwaa \times ddAa$ | 0 | $\frac{1}{2}(1-c)$ | 0 | 0 | $\frac{1}{2}(1+c)$ | 0 | 0 | 0 | 0 |
| $wwaa \times ddAA$ | 0 | 0 | 0 | 0 | 1 | 0 | 0 | 0 | 0 |
| $dwaa \times dwaa$ | $\frac{1}{4}(1-c)^2$ | $\frac{1}{2}(1-c)^2$ | $\frac{1}{4}(1-c)^2$ | $\frac{1}{2}c(1-c)$ | $c(1-c)$ | $\frac{1}{2}c(1-c)$ | $\frac{1}{4}c^2$ | $\frac{1}{2}c^2$ | $\frac{1}{4}c^2$ |
| $dwaa \times ddaa$ | 0 | $\frac{1}{2}(1-c)^2$ | $\frac{1}{2}(1-c)^2$ | 0 | $c(1-c)$ | $c(1-c)$ | 0 | $\frac{1}{2}c^2$ | $\frac{1}{2}c^2$ |
| $dwaa \times dwAa$ | $\frac{1}{8}(1-c)^2$ | $\frac{1}{4}(1-c)^2$ | $\frac{1}{8}(1-c)^2$ | $\frac{1}{8}(1-c)(1+2c)$ | $\frac{1}{4}(1-c)(1+2c)$ | $\frac{1}{8}(1-c)(1+2c)$ | $\frac{1}{8}c(1+c)$ | $\frac{1}{4}c(1+c)$ | $\frac{1}{8}c(1+c)$ |
| $dwaa \times ddAa$ | 0 | $\frac{1}{4}(1-c)^2$ | $\frac{1}{4}(1-c)^2$ | 0 | $\frac{1}{4}(1-c)(1+2c)$ | $\frac{1}{4}(1-c)(1+2c)$ | 0 | $\frac{1}{4}c(1+c)$ | $\frac{1}{4}c(1+c)$ |
| $dwaa \times ddAA$ | 0 | 0 | 0 | 0 | $\frac{1}{2}(1-c)$ | $\frac{1}{2}(1-c)$ | 0 | $\frac{1}{2}c$ | $\frac{1}{2}c$ |
| $ddaa \times ddaa^{\dagger}$ | 0 | 0 | $(1-c)^2$ | 0 | 0 | $2c(1-c)$ | 0 | 0 | $c^2$ |
| $ddaa \times dwAa$ | 0 | $\frac{1}{4}(1-c)^2$ | $\frac{1}{4}(1-c)^2$ | 0 | $\frac{1}{4}(1-c)(1+2c)$ | $\frac{1}{4}(1-c)(1+2c)$ | 0 | $\frac{1}{4}c(1+c)$ | $\frac{1}{4}c(1+c)$ |
| $ddaa \times ddAa^{\dagger}$ | 0 | 0 | $\frac{1}{2}(1-c)^2$ | 0 | 0 | $\frac{1}{2}(1-c)(1+2c)$ | 0 | 0 | $\frac{1}{2}c(1+c)$ |
| $ddaa \times ddAA^{\dagger}$ | 0 | 0 | 0 | 0 | 0 | $1-c$ | 0 | 0 | $c$ |
| $dwAa \times dwAa$ | $\frac{1}{16}(1-c)^2$ | $\frac{1}{8}(1-c)^2$ | $\frac{1}{16}(1-c)^2$ | $\frac{1}{8}(1-c)(1+c)$ | $\frac{1}{4}(1-c)(1+c)$ | $\frac{1}{8}(1-c)(1+c)$ | $\frac{1}{16}(1+c)^2$ | $\frac{1}{8}(1+c)^2$ | $\frac{1}{16}(1+c)^2$ |
| $dwAa \times ddAa$ | 0 | $\frac{1}{8}(1-c)^2$ | $\frac{1}{8}(1-c)^2$ | 0 | $\frac{1}{4}(1-c)(1+c)$ | $\frac{1}{4}(1-c)(1+c)$ | 0 | $\frac{1}{8}(1+c)^2$ | $\frac{1}{8}(1+c)^2$ |
| $dwAa \times ddAA$ | 0 | 0 | 0 | 0 | $\frac{1}{4}(1-c)$ | $\frac{1}{4}(1-c)$ | 0 | $\frac{1}{4}(1+c)$ | $\frac{1}{4}(1+c)$ |
| $ddAa \times ddAa^{\dagger}$ | 0 | 0 | $\frac{1}{4}(1-c)^2$ | 0 | 0 | $\frac{1}{2}(1-c)(1+c)$ | 0 | 0 | $\frac{1}{4}(1+c)^2$ |
| $ddAa \times ddAA^{\dagger}$ | 0 | 0 | 0 | 0 | 0 | $\frac{1}{2}(1-c)$ | 0 | 0 | $\frac{1}{2}(1+c)$ |
| $ddAA \times ddAA^{\dagger}$ | 0 | 0 | 0 | 0 | 0 | 0 | 0 | 0 | 1 |

### Supplemental Figures

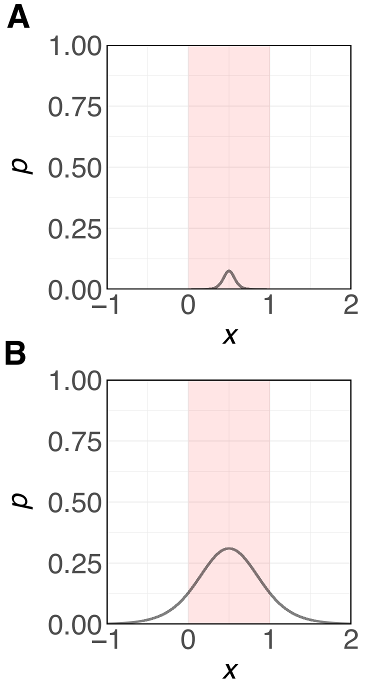


**Figure S1: Critical bubbles of two underdominance systems.** We centered the critical bubble at $x = 0.5$ in the 1D SLiM model and only modeled the portion of the bubble that fell within the SLiM boundaries (shown in red). **A.** Critical bubble of an underdominance allele with $D = 10^{-5}$, $\hat{p} = 0.05$, and $s_u = 0.05$. **B.** Critical bubble of an underdominance allele with $D = 10^{-3}$, $\hat{p} = 0.2$, and $s_u = 0.05$.

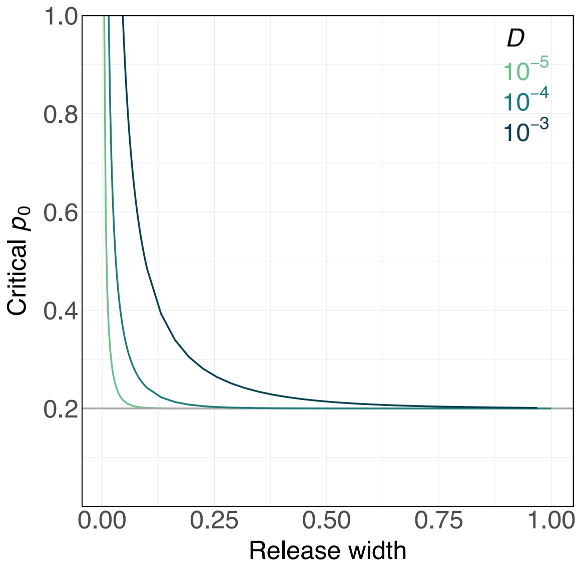


**Figure S2: Critical introduction frequencies in deSolve.** For an underdominance system with $\hat{p} = 0.2$ and $s_u = 0.8$, the critical introduction frequency was calculated for each combination of diffusion constant $D$ and release width. The critical introduction frequency is defined as the minimum introduction frequency ($p_0$) at which the underdominance allele increases in frequency in 1D reaction-diffusion simulations implemented in deSolve. The grey horizontal line marks the critical introduction frequency in a panmictic model ($\hat{p}$).

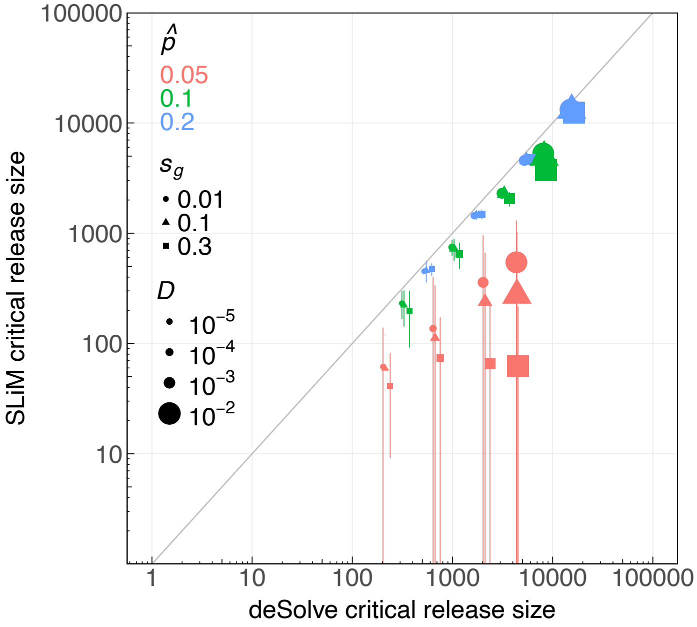


**Figure S3: Rectangular releases in SLiM versus deSolve for homing underdominance gene drives.** Drive homozygotes were introduced in a central release width in the 1D SLiM model and in 1D deSolve reaction-diffusion simulations. Each point represents a different homing underdominance gene drive, with bars indicating the drive's transition range in SLiM. We assumed an 100% germline cleavage rate and homing success rate (with no resistance). For a given fitness cost ($s_g$) and invasion threshold ($\hat{p}$), the dominance coefficient ($h$) was derived using Eq. S3.

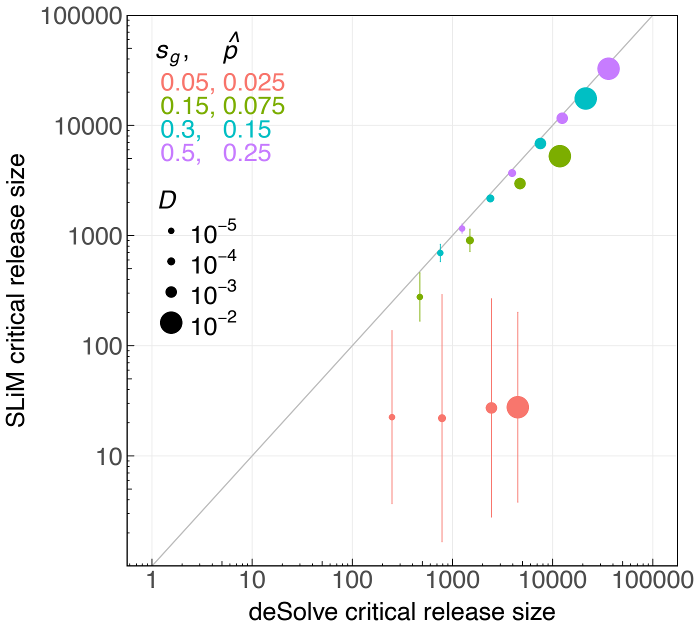


**Figure S4: Rectangular releases in SLiM versus deSolve for TADE modification drives.** This plot follows the same style as Figure 6 except depicts the results of 1D simulations, in which TADE heterozygotes were released from a central release width.

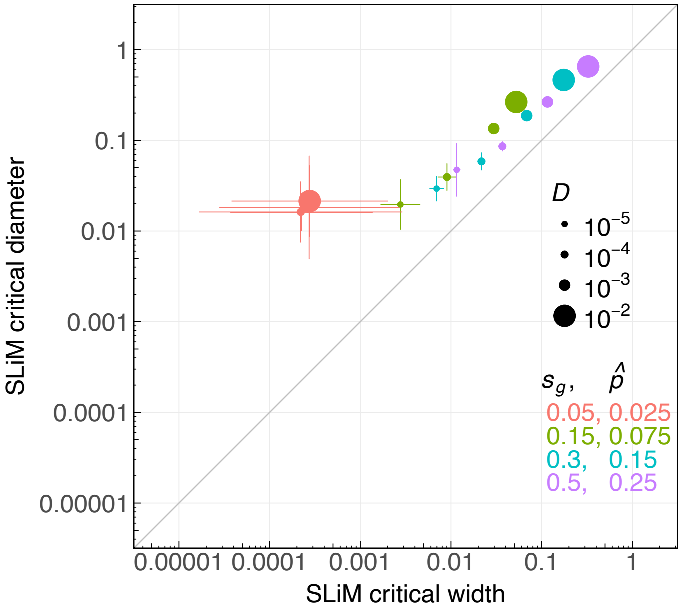


**Figure S5: SLiM critical release width (1D) versus diameter (2D) for the TADE modification drive.** TADE heterozygotes (genotype $dwAa$) were released from a central width in 1D or central circle in 2D. Each point represents a different parameterization of the drive. The $x$-axis shows the critical release width in 1D, with horizontal bars indicating the transition range in the 1D SLiM model. The $y$-axis shows the critical release diameter in 2D, with vertical bars showing the transition range in the 2D SLiM model. We set the drive dominance coefficient ($h$) to 0.5 and germline cleavage rate ($c$) to 1 such that the invasion threshold ($\hat{p}$) only depends on the drive fitness cost ($s_g$). $\hat{p}$ was found numerically based on $s_g$ (see Section **??**). The grey line denotes $y = x$.
